# Exonic enhancers are a widespread class of dual-function regulatory elements

**DOI:** 10.1101/2025.03.27.645641

**Authors:** Jean-Christophe Mouren, Magali Torres, Antoinette van Ouwerkerk, Iris Manosalva, Frederic Gallardo, Salvatore Spicuglia, Benoit Ballester

**Author notes:** Co-authors (Magali Torres, Antoinette van Ouwerkerk). Co-last authors (Salvatore Spicuglia, Benoit Ballester).

## Abstract

Exonic enhancers (EEs) occupy an under-appreciated niche in gene regulation. By integrating transcription factor binding, chromatin accessibility, and high-throughput enhancer-reporter assays, we demonstrate that many protein-coding exons possess enhancer activity across species. These EEs exhibit characteristic epigenomic signatures, form long-range interactions with gene promoters, and can be altered by both nonsynonymous and synonymous variants. CRISPR–mediated inactivation demonstrated the involvement of EEs in the cis-regulation of host and distal gene expression. Through large-scale cancer genome analyses, we reveal that EE mutations correlate with dysregulated target-gene expression and clinical outcomes, highlighting their potential relevance in disease. Evolutionary comparisons show that EEs exhibit both strong sequence constraint and lineage-specific plasticity, suggesting that they serve ancient regulatory functions while also contributing to species divergence. Our findings redefine the landscape of functional elements by establishing EEs as a component of gene regulation, while revealing how coding regions can simultaneously fulfil both protein-coding and cis-regulatory roles.

## Main

Precise control of gene expression relies on a complex network of regulatory elements that integrate spatial, temporal, and cell-type–specific signals. Traditionally, these regulatory elements have been localized to noncoding regions of the genome, where enhancers serve as key modulators of transcriptional activity^1^. However, accumulating evidence points to additional regulatory roles within protein-coding sequences, challenging the assumption that coding regions merely encode proteins^2^.

The concept of exonic enhancers (EEs) has arisen from observations that certain exons, despite encoding protein domains, also exhibit features comparable to enhancers, such as open chromatin, transcription factor (TF) occupancy, and histone modifications associated with active regulatory elements^2–4^. Early reports of enhancer-like features in certain exons suggested that this phenomenon might be more prevalent than initially assumed.

Here, we confirm that many exons act as both coding and regulatory sequences, revealing a previously underappreciated layer of genomic complexity that challenges the classic distinction between coding and noncoding regions. These dual-purpose elements can carry mutations that affect protein function and transcriptional regulation, potentially amplifying their impact on phenotypic variation and disease susceptibility.

In this study, we present a comprehensive analysis of EEs across vertebrate and invertebrate genomes using large-scale integration of TF-binding maps, open chromatin assays, and high-throughput enhancer-reporter data. We systematically identify hundreds of EEs, validate their enhancer-like properties, and explore their functional relevance in various biological contexts. By intersecting EEs with cancer genomic datasets, we demonstrate that variants within these coding-regulatory regions can influence gene expression and patient outcomes, suggesting broad clinical implications. Finally, through multi-species comparative analyses, we reveal that EEs display both deep evolutionary conservation and lineage-specific diversification, reflecting selective pressures to maintain dual coding-enhancer functions while enabling adaptive regulatory plasticity.

Collectively, our findings establish EEs as an integral, albeit underrecognized, layer of gene regulation. They emphasize the need to revisit the traditional paradigm that confines enhancers strictly to noncoding DNA, highlighting how coding exons themselves can shape gene expression in both health and disease. This broader perspective on exonic functionality has important implications for variant interpretation, disease mechanisms, and the overall architecture of the genome.

## Results

### Exonic Enhancers in four species revealed by TF binding maps

To investigate whether protein-coding exons can harbor regulatory activity, we integrated large-scale TF ChIP-seq profiles from ReMap^5^ to construct a high-resolution map of TF occupancy in coding sequences. We observed a pronounced enrichment of multi–TF binding in certain internal exons, as illustrated by the GARRE1 and USP20 loci, which exhibit representative regulatory signatures (Fig. 1a). In line with previous reports of DNase I hypersensitive sites in exonic regions^2^, we found that 3.9% of DNase I hypersensitive sites overlapped exonic regions, underscoring the regulatory potential of exons (Fig. 1b). In parallel, TF ChIP-seq peaks occupy diverse genomic contexts, with the majority of ReMap peaks mapping to intronic (40.9%) and intergenic (25.6%) regions—consistent with traditional enhancers—yet 4.2% of these peaks localise to coding exons (Fig. 1b). Moreover, exonic regions overlapping classic regulatory features—such as enhancer marks from ENCODE cCREs^6^, DNase I hypersensitive sites^7^, ATAC-seq peaks (ChIP-Atlas^8^), CAGE TSS (FANTOM^9,10^), and histone modifications like H3K27ac^6^ (Fig. 1c)—further reinforce the notion that some exons could actively participate in transcriptional regulation.

**Figure 1.**
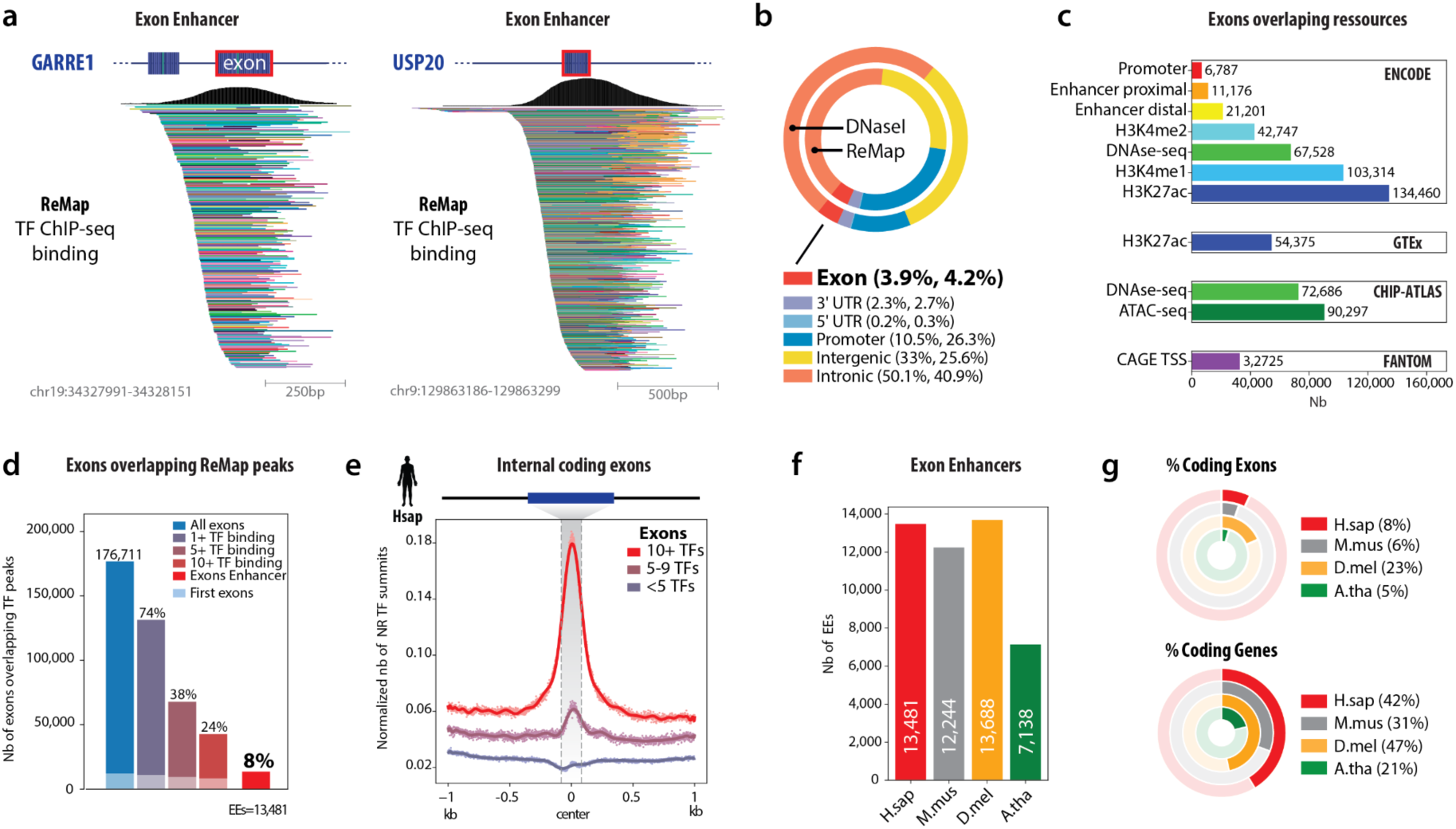
Identification and characterization of Exon Enhancers (EEs) across species. (**a**) Two representative examples of EEs within protein-coding exons (GARRE1 on human chromosome 19 and USP20 on chromosome 9). The top black traces depict aggregated TF ChIP-seq signal (ReMap), illustrating a peak of transcription factor (TF) occupancy centered on an internal exon. Each coloured bar below represents a distinct TF dataset from ReMap, collectively underscoring extensive multi-TF binding at these exons. (**b**) Donut charts showing the proportion of genomic annotations (e.g., intronic, intergenic, promoter, UTR, exon) identified by DNase I hypersensitivity (outer ring) and TF occupancy from ReMap (inner ring). Notably, exons account for ∼4% of these regulatory regions, illustrating that a non-negligible fraction of regulatory elements localizes to protein-coding exons. (**c**) Bar plots indicating the overlap of exons with various genomic resources, including ENCODE (e.g., promoter, enhancer, DNAse-seq, histones marks), GTEx (H3K27ac histone mark), CHiP-ATLAS (DNAse-seq, ATAC-seq), and FANTOM (CAGE TSS). A substantial number of exons intersect enhancer-related annotations, implying that coding exons frequently harbour regulatory potential. (**d**) Histogram depicting the total number of exons overlapping TF-bound peaks according to ReMap, stratified by the minimum number of TFs bound. Of 176,711 exons, 74% have ≥1 TF, 38% have ≥5 TFs, 24% have ≥10 TFs, and 8% are designated as “exon enhancers” (EEs). This suggests that EEs, although a minority, are enriched for multiple overlapping TF-binding events. (**e**) Metagene plot of TF summit density centered on internal coding exons. Exons bound by ≥10 TFs (red trace) exhibit a strong, focused peak at the exon center compared to exons bound by 5–10 TFs (purple trace) or <5 TFs (grey trace), indicating that highly TF-occupied exons are enriched in regulatory activity precisely at their midpoint. (**f**) Bar chart showing the number of EEs detected in four model organisms—human (H. sapiens), mouse (M. musculus), fruit fly (D. melanogaster), and thale cress (A. thaliana). The presence of EEs across diverse species underscores the evolutionary breadth of exon-centered regulatory elements. (g) Donut plots summarizing the proportion of coding exons (top) and coding genes (bottom) in each species. Although D. melanogaster has a higher fraction of coding exons (23%) compared to the other species examined, the percentage of EEs is nonetheless appreciable in all four genomes, suggesting a broadly conserved role for EEs in gene regulation.

Across the human genome, 74% of coding exons (n=131,358) overlap at least one TF-binding peak, and 24% were bound by ten or more TFs (n=42,699), highlighting the frequent occurrence of TF occupancy within coding exons (Fig. 1d). Notably, 8% of these TF-bound exons (n=13,481) satisfy stringent multi–TF binding thresholds and filtering methods, displaying enhancer-like activity. We therefore designated these 13,481 exons as exonic enhancers (EEs) in the rest of this study. We identified EEs using stringent criteria to distinguish them from promoters or other regulatory elements while ensuring TF binding was predominantly confined to the exon, which prevented confounding signals from adjacent intronic regions (Methods). Consequently, some previously tested EEs from the literature^4^ were excluded (Supplementary Fig. 1). Meta-exon analyses show a focused peak of TF summits at the midpoint of these exons, particularly among those with extensive TF co-occupancy (Fig. 1e). To evaluate EEs beyond human, we extended our TF overlap analysis to mouse (*Mus musculus*), fruit fly (*Drosophila melanogaster*), and thale cress (*Arabidopsis thaliana*) (Supplementary Fig. 2-5). In all four species, coding exons show evidence of enhancer-like features, including TF occupancy (Supplementary Fig. 3-6) and chromatin accessibility (Supplementary Fig. 2), indicating that EEs are evolutionarily widespread (Fig. 1f, Mmus EEs=12,244; Dmel EEs=13,688; Atha EEs=7,138). Although the fraction of exons and genes differs among these organisms (Fig. 1g), a consistent presence of EEs suggests that these elements represent a conserved facet of gene regulation across species (Supplementary Fig. 7). Our cross-species analyses establish EEs as a fundamental yet previously under-appreciated component of the regulatory landscape, revealing a substantial layer of complexity within coding exons and underscoring their pivotal role in gene regulation.

### Exonic Enhancers display enhancer-associated features across genomes

To assess whether EEs display canonical enhancer-associated features, we integrated publicly available DNase I hypersensitivity and ATAC-seq datasets from ENCODE^6^ and ChIP-Atlas^8^. Across the four species, we observed a high degree of overlap between EEs and open chromatin marks (Fig. 2a, Supplementary Fig. 8). Notably, 82% of human EEs and 67% of mouse EEs overlapped with DNase or ATAC peaks, and we also detected strong concordance with H3K27ac and H3K4me1,2 histone marks (Supplementary Fig. 9), suggesting that EEs adopt active chromatin configurations conducive to TF binding. These observations echo earlier findings that DNase I hypersensitive and ATAC-accessible sites often reflect enhancer potential^11^.

**Figure 2.**
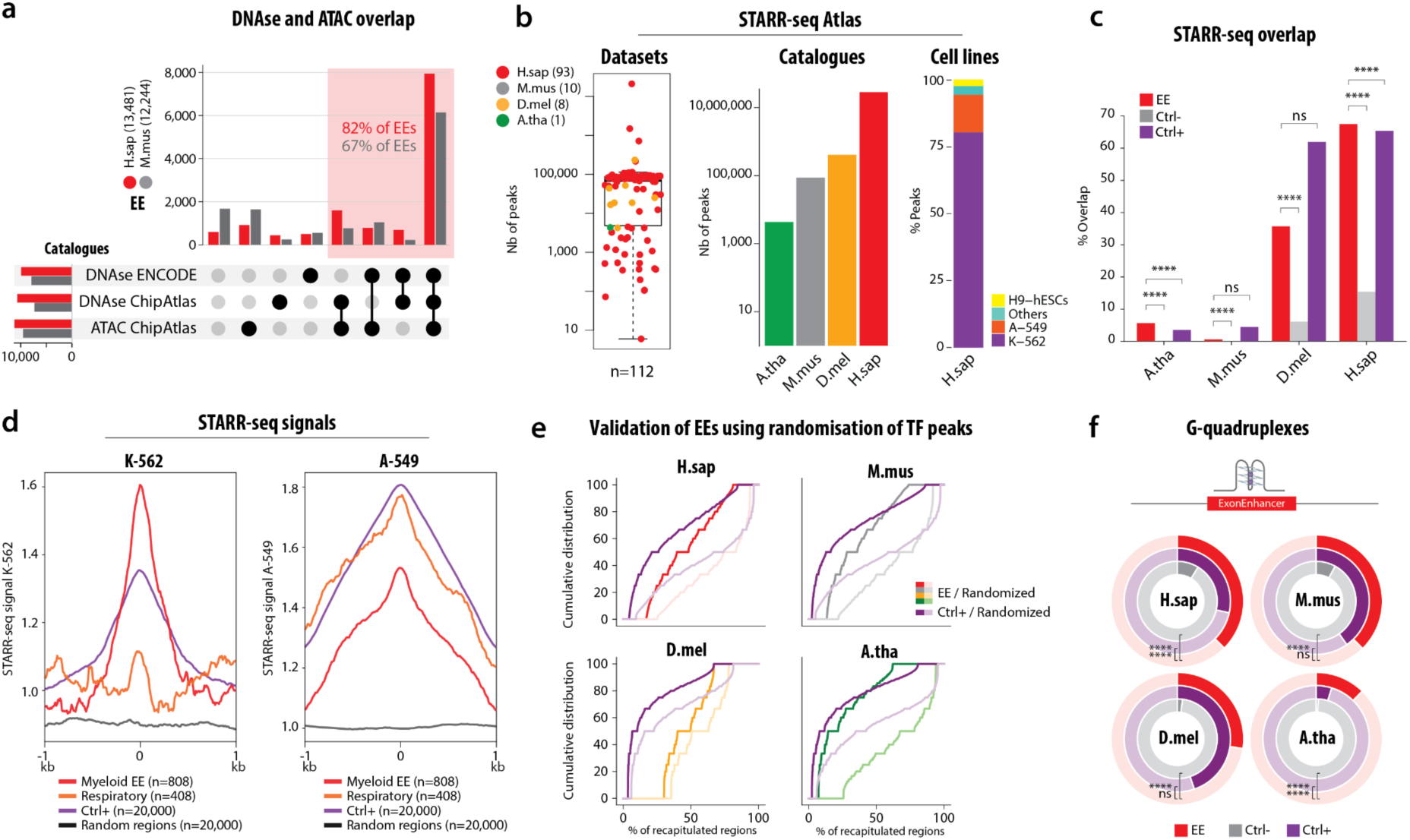
Exonic Enhancers display enhancer-associated features across genomes. **(a)** Combinatorial overlaps of EEs with DNase I hypersensitive sites and ATAC-seq datasets, derived from ENCODE and ChIP-Atlas resources, for *H. sapiens* (H.sap) and *M. musculus* (M.mus). The upset plot highlights that 82% of human EEs and 67% of mouse EEs exhibit open chromatin signals (red shading). **(b)** STARR-seq Atlas summary. The box-and-whisker plot (left) illustrates the number of STARR-seq peaks identified per dataset (n=112 total datasets), color-coded by species. The middle bar graph depicts the total count of STARR-seq peaks per species, and the stacked bar chart (right) shows cell line or tissue composition in the human dataset (e.g., K-562, A-549, H9-hESCs). **(c)** Comparison of STARR-seq overlap percentages for EEs (red), negative controls (Ctrl–, grey), and positive controls (Ctrl+, purple) across the four species. Statistical significance (Fisher’s exact one-side test) is indicated: ****P < 0.00001, ns = not significant. **(d)** Metaprofiles depicting STARR-seq signal intensity in human K-562 (left) and A-549 (right) cells, centered on exon enhancers (EEs) identified with myeloid (red) and respiratory (orange) signatures. Signals from positive control regions (Ctrl+, purple), negative controls (random genomic regions; dotted grey line) are shown for comparison. Signal enrichment demonstrates significant enhancer activity specifically around tissue-associated Ees **(e)** Validation of EEs using randomization of TF peaks. Cumulative distribution curves compare the fraction of EEs (red) and positive controls (purple) against randomized TF peaks (light lines) for each species, indicating that EEs are significantly enriched for TF binding relative to random expectation. **(f)** G-quadruplex (G4) content within EEs (red arcs), negative controls (grey arcs), and positive controls (purple arcs) across *H. sapiens*, *M. musculus*, *D. melanogaster*, and *A. thaliana*. The doughnut plots highlight significantly higher G4 occurrences in EEs relative to controls (****P < 0.00001, ns = not significant, Fisher exact one-side test).

To functionally validate the enhancer-like activity of EEs, we compiled a comprehensive multi-species STARR-seq atlas spanning 112 datasets across multiple cell lines and tissues (Fig. 2b, Methods, Supp File). STARR-seq, which quantifies enhancer-driven transcription in a plasmid-based reporter assay^12^, revealed millions of potential enhancer peaks, including a discrete subset within EEs. Across all four species, EEs exhibited significantly higher overlap with STARR-seq peaks than coding exons lacking TF peaks (Ctrl-) (Fig. 2c) and overlap rates comparable to classic enhancers (Ctrl+), defined as intergenic distal cCREs overlapping at least 10 TFs and selected using the same EE filtering criteria (Methods), suggesting that EEs are more likely to drive reporter gene expression. Furthermore, metaprofile analyses of STARR-seq data from K-562 (erythromyeloid) and A-549 (respiratory epithelial) cells revealed tissue-specific elevated enhancer signals, notably enriched at exon enhancers associated with myeloid and respiratory tissues, respectively, compared to control and random genomic regions (Fig. 2d, Supplementary Fig. 10, Methods).

To validate the regulatory potential of EEs computationally, we hypothesized that the ChIP-seq signals detected in EEs might be supported by an enrichment of transcription factor binding sites (TFBS) aligning with the TF peaks, similar to classic enhancers (Ctrl+) (Fig. 2e). Using a genome-wide randomization strategy, we shuffled TF peaks across each genome assembly and assessed binding motif occupancy. Remarkably, EEs maintained a significantly higher occupancy of matching TF with their corresponding binding sites across all four species, compared to randomized conditions. This pattern, combined with a higher TFBS density in EEs (Supplementary Fig. 11), mirrors the behaviour of classical enhancers and exceeds expectations based on random distributions. These findings establish EEs as a distinct and functional class of enhancers, rather than mere artifacts arising from proximal ligation during chromatin immunoprecipitation, contrasting with earlier interpretations^13^. They also align with the dual role of protein-coding exons as TF-binding platforms^2^ thereby extending the concept of exon-centered regulation.

Previous studies implicated secondary DNA structures in enhancer function^14^. We analysed G-quadruplex (G4) motifs in EEs versus matched controls (Ctrl– and Ctrl+) across four species. EEs showed significantly higher predicted G4 sequences and GC content (Fig. 2f, Supplementary Fig. 12), suggesting that G4 formation may enhance TF binding or modulate transcription^15^. In conclusion, our multi-species analyses reveal that many exons act as bona fide enhancers, characterized by open chromatin, STARR-seq activity, relevant TF binding sites, and an enrichment of G4 and GC-rich sequences.

### Functional validation of Exonic Enhancers activity

To experimentally confirm the regulatory potential of EEs, we selected a panel of candidates identified in our integrative analysis (Fig. 1 and 2) and cloned them into luciferase reporter constructs (Fig. 3a). Each EE was inserted upstream of a minimal SV40 promoter, allowing quantification of enhancer-driven transcription. In total, we tested 20 predicted EEs in luciferase reporter assays (Fig. 3a). Of these, 8 exhibited moderate to robust transcriptional activation (fold-change range 1.6–11.5) compared to both empty vector and exon-derived control sequences (2 exons without TF, 1 EE previously identified in HepG2^4^. These results align with enhancer-like profiles inferred from histone modifications (H3K27ac, H3K4me1), TF ChIP-seq occupancy, and chromatin accessibility (DNase/ATAC-seq).

**Figure 3.**
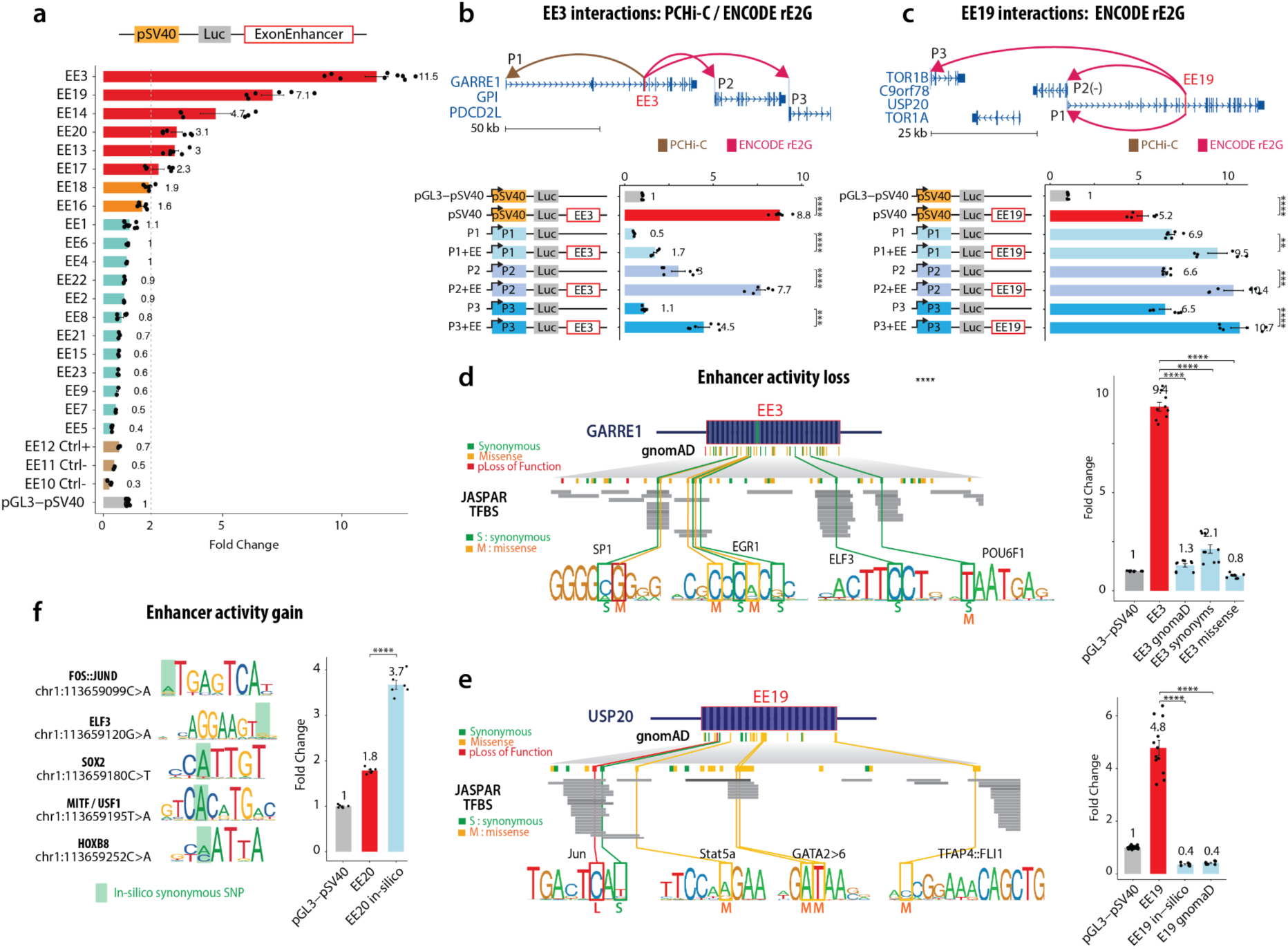
Functional characterization of EEs and the impact of genetic variants. **(a)** Luciferase reporter assays for a panel of exonic enhancers (EEs) cloned upstream of the SV40 promoter. Bars represent the relative fold-change in luciferase activity compared to the empty pGL3– pSV40 vector, with EEs showing differential enhancer activity. Control constructs are indicated (Ctrl). **(b)** PCHi-C and ENCODE-rE2G analyses reveal physical interactions between EE3 and nearby gene promoters (*GARR1, GPI, PDCD2L*). The schematic (top) depicts chromatin looping (arcs) linking EE3 to three putative promoters (P1–P3). The bar chart (bottom) shows luciferase activities of combinations of EEs and promoters, indicating synergistic enhancer effects. ****P < 0.00001 t-test. **(c)** Similarly, EE19 engages promoters of *TOR1B, C9orf78, USP20,* and *TOR1A* through long-range chromatin interactions (top). Luciferase assays (bottom) recapitulate these contact-based enhancer-promoter relationships, further supporting the regulatory potency of EE19. Error bars in all panels represent mean ± s.e.m. for at least three biological replicates. ****P < 0.00001 t-test. **(d,e)** Representative examples of genetic variants affecting EE function in the *GARRE1* (EE3) and *USP20* (EE19) loci, respectively. The schematics depict gnomAD variant classes—synonymous (S), missense (M), and loss-of-function (L)—along with predicted transcription factor (TF) binding sites based on JASPAR motifs. The bar plots at right show the resulting luciferase activity of reference and variant EE constructs relative to pGL3–pSV40. Constructs either have only missenses, synonymous or a mix of the several variant classes. EE19 *in-silico* construct consists of synonymous mutations. ****P < 0.00001 t-test. **(f)** Variants that confer enhancer activity gain in selected TF binding sites. Sequence logos illustrate key TF motifs (e.g., FOS::JUND, EBF3, SOX9, MTF1/USF1, HOXB), with *in silico* synonymous SNPs highlighted in green. Fold-change data compare EE-driven reporter activity against pGL3–pSV40. ****P < 0.00001 t-test.

To investigate the effect of selected EEs (e.g., EE3 and EE19) on promoters of host and distal genes, we integrated promoter capture Hi-C (PCHi-C)^16^ and ENCODE-rE2G^17^ datasets (Fig. 3b,c). These analyses revealed physical contacts spanning tens of kilobases between each EE and its host gene promoter, as well as distal promoters, suggesting a long-range regulatory relationship. To validate this, we replaced the standard SV40 promoter in our luciferase constructs with the predicted target promoters (P1, P2, P3) of two of our strongest EEs (EE3 and EE19). Reporter assays demonstrated a significant increase in activity for all three EE3 and EE19 associated promoters, supporting the notion that these EEs can engage multiple promoters across substantial genomic distances. These findings indicate that EEs can directly engage their endogenous promoters over substantial genomic distances, underscoring their role in long-range gene regulation.

### Coding variants influencing Exonic Enhancer function

We next investigated how naturally occurring genetic variation affects EE activity. Using variant data from gnomAD^18^, we focused on synonymous, missense, and loss-of-function substitutions that overlapped predicted TFBS. At two representative loci, *GARRE1* (EE3) and *USP20* (EE19), we detected synonymous and missense variants within TF consensus sites, as predicted by JASPAR^19^. Reporter luciferase assays revealed that these variants disrupted enhancer activity, leading to a significant reduction or complete loss of function (Fig. 3d,e).

Conversely, *in silico* introduction of synonymous variants designed to create or strengthen TF-binding motifs within the EE20 sequence produced an increase in luciferase signal (Fig. 3f). Hence, even ostensibly ‘silent’ mutations in coding regions can have a regulatory impact by altering enhancer function. By coupling mutational data with functional assays, we demonstrate that EEs can acquire gain- or loss-of-function phenotypes in the presence of coding mutations. Collectively, these observations position EEs as not only important cis-regulatory elements in normal gene expression but also as potential contributors to pathological states when harbouring disruptive variants.

### STARR-seq validates hundreds of Exonic Enhancers

We previously observed that many EEs overlapped with publicly available STARR-seq predictions, yet those predicted enhancers often included adjacent intronic regions, obscuring whether the exonic sequence alone was responsible for enhancer activity. To unambiguously determine the genome-wide enhancer potential of EEs, we performed STARR-seq assays in K-562 cells using a comprehensive library of exonic fragments (n=2,068, excluding surrounding intronic sequences), alongside positive and negative control sequences (Fig. 4a, Methods). This assay positions EE candidate elements downstream of a minimal promoter, enabling direct and quantitative measurement of enhancer-driven transcription^12^. Across replicates, K-562 EEs exhibited significantly higher STARR-seq activity than negative controls (Ctrl–, *n*=1,235, p=8.737×10^-20^ t-test) and, in many cases, approached or exceeded the activity levels of established classic enhancers (Ctrl+, *n*=1,184, p=1,650×10^-22^ t-test) (Fig. 4b). EEs associated with A-549 or GM12878 cell line signatures show significantly lower activity in K-562 cells, suggesting context-dependent regulatory function (Supplementary Fig. 13). These findings reveal that a substantial fraction of coding DNA harbours strong enhancer activity, further highlighting its regulatory potential.

**Figure 4:**
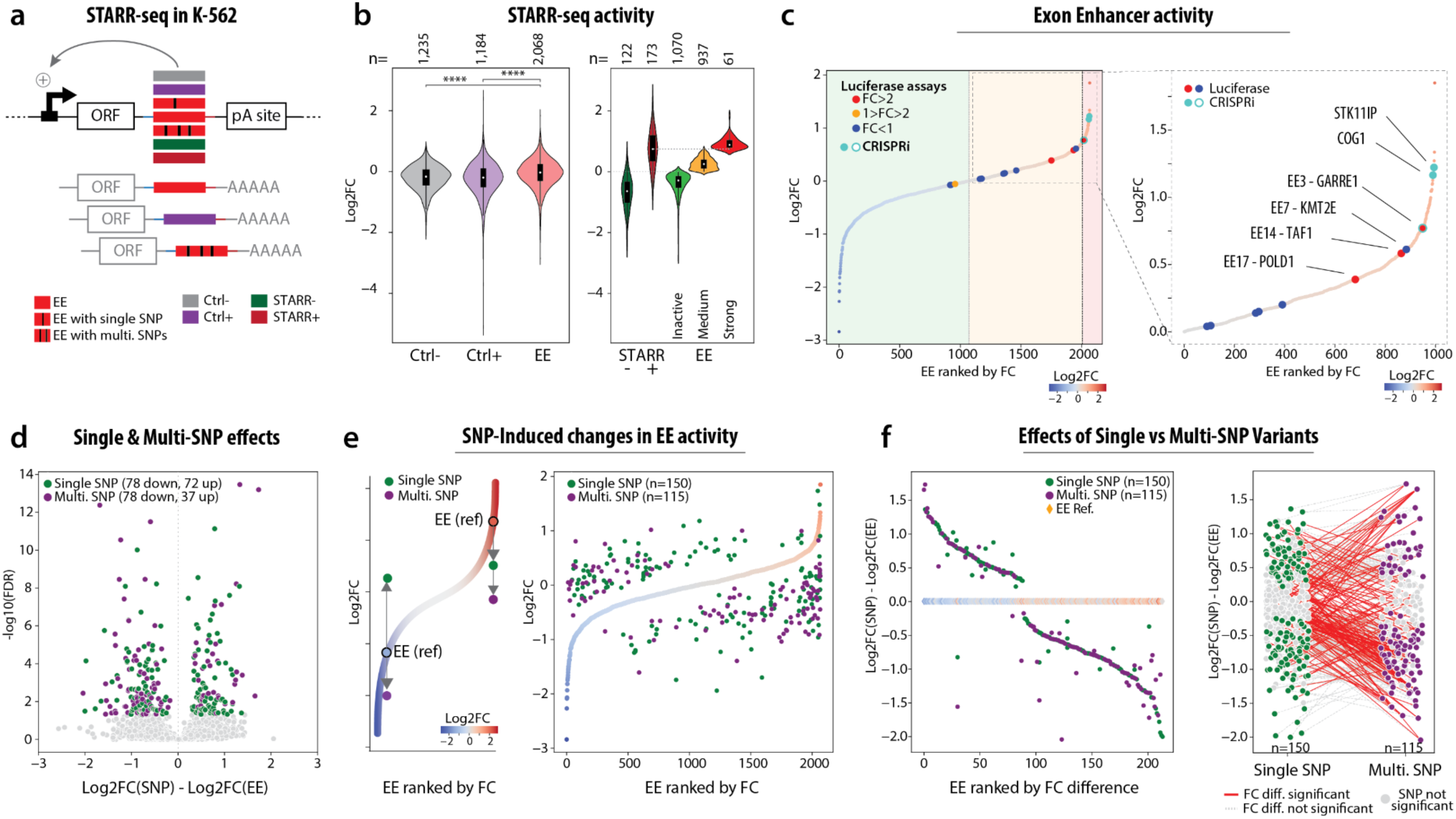
STARR-seq analysis of EEs and the impact of single and multiple SNPs on enhancer activity. **(a)** Schematic overview of the STARR-seq experimental design in K-562 cells. Exon Enhancers (EEs; red) and control sequences (Ctrl–, purple; Ctrl+, grey, STARR+ dark red, STARR-dark green) were cloned downstream of a constitutively active promoter driving a reporter ORF and upstream of a poly(A) signal. Both synonymous single-SNP and multi-SNP variants were assessed. **(b)** Violin plots showing the distribution of STARR-seq activity (log2 fold change) for negative controls (Ctrl–), positive controls (Ctrl+), and EEs. STARR± indicates previously identified regions with weak and strong STARR-seq activity. Statistical significance is denoted: ****P < 0.00001 Student’s t-test. **(c)** Rank-ordered plot of EEs by STARR-seq fold change, with color-coded luciferase outcomes (red, orange, blue). The right-hand zoom highlights top-ranked EEs and EEs tested by luciferase reporter assays labeled by the associated gene locus (e.g., MAP3K13, SERTAD2, EE3–GARRE1, EE7– KMT2E, EE14–TAF1, EE17–POLD1). **(d)** Scatterplot of variant effects. Single-SNP (green) and multi-SNP (purple) EEs are plotted based on the difference between log2 fold change of the reference EE and variant EE on the x-axis, and – log10(FDR) (Padj < 0.05, Benjamini and Hochberg), on the y-axis. Points above the grey background indicate significant divergence in enhancer activity linked to SNP presence. **(e)** Right panel: rank-ordered distribution of EEs by STARR-seq fold change. Reference EEs are contrasted with variant EEs carrying single (green) or multiple (purple) SNPs. **(f)** Left panel: difference in log2 fold change (EE[SNP] – EE) is plotted for single- and multi-SNP subsets, indicating the extent to which variant alleles modify enhancer activity. Right panel: Comparison of STARR-seq activity changes for single-vs. multi-SNP EEs, shown as log2 fold-change (EE harbouring SNP - EE reference) on the y-axis. Green circles denote EEs with single-SNP variants (n=150), purple circles denote EE with multi-SNP variants (n=115). Connecting lines link EEs harbouring both single- and multi-SNP variants; red lines indicate significant differences (Padj < 0.05, Benjamini and Hochberg), and grey lines indicate nonsignificant differences.

To define thresholds for EE activity, we used controls, promoter elements previously characterized with enhancer-like activity (STARR+, n=173), and their inactive counterparts (STARR–, n=122), validated through prior STARR-seq studies^20^. Based on these benchmarks, we categorized EEs into three groups: inactive EEs (green, fold change [FC] ≤ 0), moderately active EEs (orange, 0 < FC < median STARR+), and highly active EEs (red, FC ≥ median STARR+) (Fig. 4b). Notably, 9 EEs validated via luciferase assays (Fig. 3a) exhibited STARR-seq signals which are globally consistent with these activity thresholds (Fig. 4c). Among the top-tier EEs, we identified loci linked to essential cellular processes, including signal transduction (MAP3K13), transcriptional regulation (TAF1), and DNA replication (POLD1) (Fig. 4c). These results establish EEs as key cis-regulatory elements capable of driving robust transcriptional activity within coding regions.

### Coding variants modulate Exonic Enhancers activity

To assess whether natural genetic variation can modulate EE activity, we introduced single-nucleotide polymorphisms (SNPs) from gnomAD^18^ predicted to alter TFBS, into our EE STARR-seq library (Fig. 4a). We tested single-SNP EEs with synonymous variants as well as multi-SNPs EEs with diverse functional impacts. STARR-seq measurements revealed that some variant EEs deviated from their reference sequence activity, indicating that coding-region mutations can alter enhancer output (Fig. 4d,e). The volcano plot (Fig. 4d) shows that EEs carrying multiple SNPs often exhibit a reduced enhancer activity (n=78 down, n=37 up), presumably due to the cumulative disruption of TF-binding motifs or other regulatory features. Meanwhile, rank-ordered analyses (Fig. 4e) confirm that both single- and multi-SNP EE variants diverge from the EE reference activity to varying extents, suggesting that coding-region variations can influence enhancer activity, as demonstrated at a lower scale (Fig3. b-d). Strikingly, variants in strong EEs tend to decrease enhancer activity while those in weak or inactive EEs tend to increase enhancer activity. Likewise, comparisons of log₂ fold-change between reference EEs and their mutated counterparts (single- or multi-SNP) illustrate a broad spectrum of activity alterations (Fig. 4f, right). In particular, multi-SNP variants show more pronounced deviations from the reference enhancer activity than single-SNP variants (Fig. 4f, left), emphasizing how multiple concurrent mutations can amplify changes in enhancer function. Combining high-throughput STARR-seq with targeted SNP mutagenesis, we demonstrate that while EEs drive robust transcriptional activity, they are remarkably sensitive to even minor coding-region variants—underscoring both their role as essential cis-regulatory platforms and their vulnerability to mutations that may lead to functional or pathological consequences.

### Exonic Enhancers form robust enhancers–target gene interactions

Having established that EEs can function as *bona fide* enhancers using STARR-seq (Fig. 4), we next sought to identify their potential target genes. To this end, we generated an interaction atlas by integrating three complementary datasets: (i) a compiled promoter capture Hi-C collection^16^, (ii) enhancer–promoter regulatory interactions from the ENCODE-rE2G resource^17^ and (iii) eQTL associations from GTEx^21^. We focused on overlap between EEs and these interaction signals, comparing them with both classic enhancers (Ctrl+) and normal coding exons (Ctrl-). Overall, EEs exhibited broader and more frequent overlaps with at least one of the three datasets, comparable to classic enhancers and substantially exceeding the negative controls even at cellular or tissue contexts (Fig. 5a, Supplementary Fig. 14). We categorized EE–target gene interactions according to whether the target gene was the EE host gene (internal), located elsewhere in the genome (external), or both (mixed). To increase the reliability of these associations for subsequent analyses, we restricted analyses to robust EE–target gene pairs (EE–TG) supported by at least two of the three resources (promoter capture Hi-C, ENCODE-rE2G, or GTEx eQTL). Notably, 26% of EEs showed such robust interactions, compared with 35% for classic enhancers and only 2% for negative controls (Fig. 5b).

**Figure 5.**
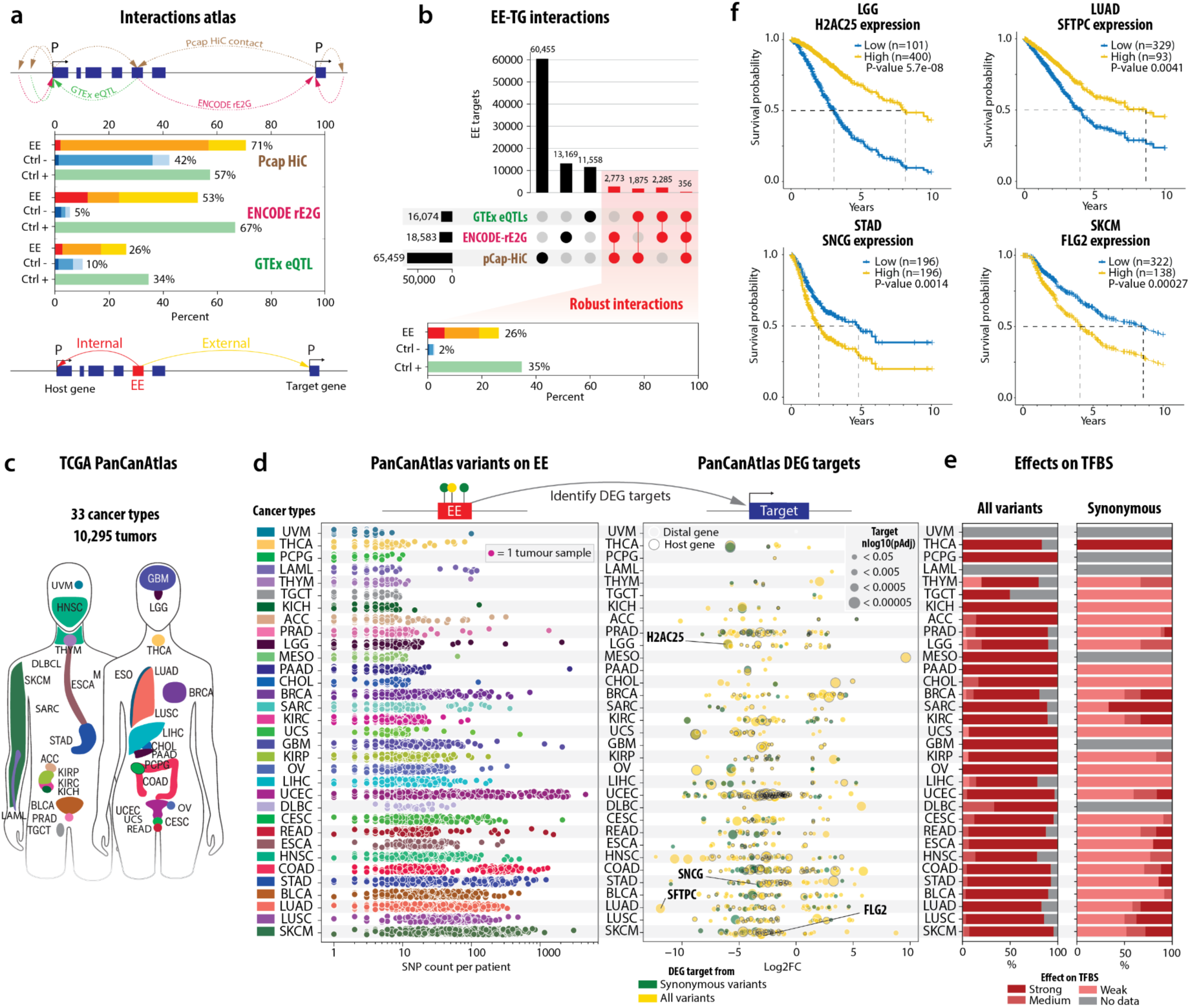
Pan-cancer analysis of Exonic Enhancers (EEs) and their target gene interactions. **(a)** Construction of an “interaction atlas” to map potential EE target genes, drawing on promoter capture Hi-C data from Laverré *et al.* (1), ENCODE-rE2G from Gschwind *et al.* (2), and GTEx eQTLs (3). The schematic shows internal (red), external (yellow), and mixed (orange) EE–target gene configurations, with bar plots indicating that EEs exhibit robust overlaps in these data sets, similar to positive controls (Ctrl+, green) and exceeding negative controls (Ctrl–, blue). **(b)** For downstream analyses, only EE–target gene (EE–TG) interactions confirmed by at least two of the three resources were retained, yielding 26% robust interactions for EEs, compared to 35% for Ctrl+ enhancers and 2% for Ctrl– sequences. **(c)** Overview of the TCGA PanCanAtlas dataset—encompassing 33 cancer types (∼10,000 tumours)— used to explore possible pathogenic effects of coding variants within EEs. Cancer sites are colour-labelled based on prior nomenclature. **(d)** Left panel: distribution of PanCancer variants per patient located within identified EEs across various tumour types. Right panel: corresponding differentially expressed target genes (DEGs) identified through EE–TG interactions. Each row represents a "flattened volcano" plot, illustrating expression changes in target genes, with DEGs linked to synonymous variants shown in green and those linked to all variants shown in yellow. **(e)** Predicted impact of PanCancerAtlas variants (left) and synonymous variants (right) on TFBSs within EEs. Disruptive effects were classified as strong (dark red), medium (red), or weak (light red) based on FABIAN-predicted binding disruption scores using JASPAR TFBS motifs. Gray bars indicate cancer types with no available data, or non-computed disruption score due to the variant type. **(f)** Survival curves showing differential overall survival (OS) based on the expression levels of four example EE-regulated target genes (*H2AC2S*, *SFTPC*, *SNCG*, and *FLG2*) in LGG, LUAD, STAD, and SKCM, respectively. The analysis was performed using a Cox proportional-hazards (PH) model. High-expression (yellow) and low-expression (blue) cohorts demonstrate significant survival differences, highlighting the potential prognostic value of these EE target genes.

These findings underscore that EEs not only display enhancer-like regulatory potential (Fig. 1-4) but also form stable interactions with downstream targets.

### Genetic variation in Exonic Enhancers impact cancer-related expression

We next addressed whether variants in EEs could have pathological or functional outcomes in human cancers. To capture this variation at scale, we utilised the TCGA Pan Cancer Atlas^22^ comprising over 10,000 tumour samples across 33 cancer types (Fig. 5c). To evaluate whether EE variants influence the expression of their putative target genes in a pathological context, we intersected the PanCanAtlas variants located in EEs with the robust EE–TG pairs previously defined (Fig. 5d). Across the 33 tumour types, the number of EE variants per patient ranged from a single event to thousands, highlighting the broad heterogeneity of mutational burden in cancer (Fig. 5d left). For each variant category—all variants (yellow) and synonymous variants only (green)—we performed differential expression analyses of the corresponding target genes, visualized as flattened volcano plots (Fig. 5d right). Notably, a subset of EE–target gene pairs exhibited significant transcriptional changes, suggesting that EE mutations may disrupt cis-regulatory architecture to drive oncogenic or impair tumour-suppressive pathways. Importantly, even synonymous substitutions (green)—historically deemed ‘silent’—were associated with altered expression (Fig. 5d right), demonstrating that such variants can subtly modulate enhancer activity. Across cancer types, TCGA variants intersecting EE frequently altered TF binding (Fig. 5e), with the effects ranging from weak (light red) to strong (dark red). Notably, synonymous mutations alone also impacted TFBSs (Fig. 5e, right), reinforcing their regulatory significance independent of coding changes. To illustrate our prediction of TF binding alterations for EEs (Supplementary Fig. 15), we showcase an EE of the MMRN2 gene (Supplementary Fig. 16), with predicted changes in TF binding affinity visualized via a public UCSC trackhub (Methods). Collectively, these findings indicate that EEs are susceptible to pathological rewiring in cancer, potentially influencing disease progression through the dysregulation of critical target genes.

### Prognostic relevance of Exonic Enhancers–target gene disruption

To evaluate whether EE-mediated regulatory changes may have clinical relevance, we selected examples of differentially expressed target genes identified in four cancer types: lower-grade glioma (LGG), lung adenocarcinoma (LUAD), stomach adenocarcinoma (STAD), and skin cutaneous melanoma (SKCM). For each of them, we assessed the prognostic value of high or low target-gene expression (Fig. 5f). In all four cancers, altered expression correlated with differential overall survival, signifying that variants located within EEs can not only perturb enhancer activity but may also drive meaningful shifts in tumour cell biology. Although these correlations suggest a role for EE disruptions in tumorigenesis, direct in vivo validation lies beyond our scope, so the link remains primarily inferential. Additionally, EEs harbour a higher fraction of disease- and measurement-related GWAS variants than controls and are enriched for pleiotropic SNPs that exhibit robust regulatory interactions (Supplementary Fig. 17). These results emphasize that exonic enhancer elements represent a class of regulatory sites highly sensitive to natural and pathological DNA variation, with potential consequences for cancer development and patient outcomes.

### CRISPRi silencing validates Exonic Enhancers-mediated gene regulation

To assess whether EEs actively modulate gene expression in vivo, we employed a potent CRISPR interference system (dCas9–KRAB–MeCP2) targeting four selected EEs in K-562 cells (Fig. 6a). Each EE was previously implicated as a regulatory hub in our interaction catalogue, contacting one or more host or distal target genes via promoter capture Hi-C, ENCODE-rE2G and GTEx eQTL mapping. We monitored the effects of EE silencing on the expression of the host (placing primers upstream and downstream of each EE to capture potentially truncated transcripts) and neighbouring genes by RT– qPCR. In all four cases we observed a regulatory effect of the EE inactivation. Notably, repression of EE3 GARRE1 and EE STK11IP resulted in significant reduction of the host and one of the distal interacting genes (GPI and CHPF, respectively) (Fig. 6b,d), while repression of EE19 USP20 and EE COG1 resulted in significant reduction of the host gene only. (Fig. 6c,e). These results demonstrate that EEs function as bona fide regulatory elements, contributing to long-range control of gene expression in the mammalian genome.

**Figure 6.**
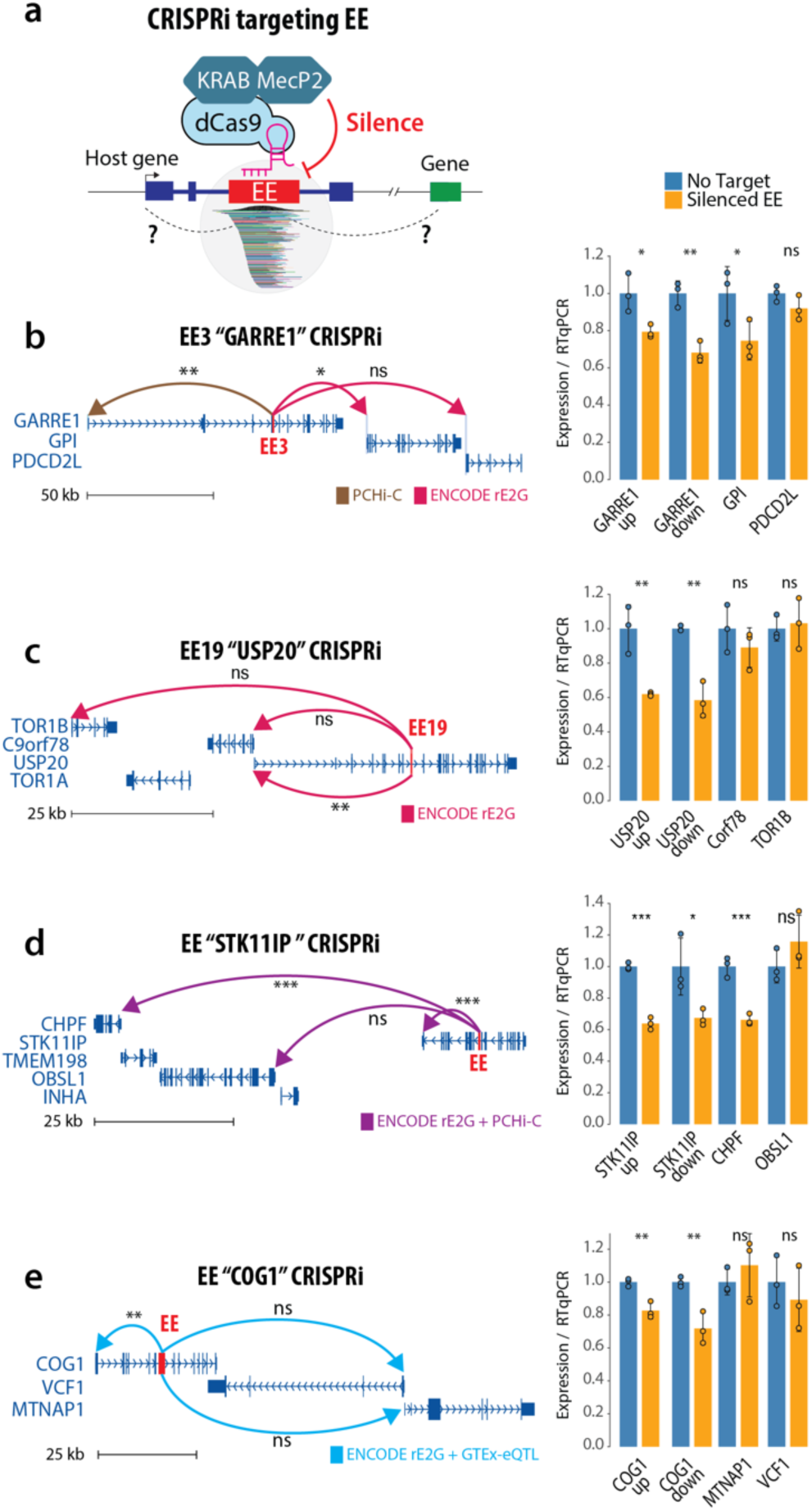
CRISPRi silencing validates EE-mediated gene regulation. (**a**) Schematic of the CRISPR interference approach using dCas9–KRAB–MeCP2 targeted to exonic enhancers (EEs). (**b–e**) Four representative EEs (EE3 GARRE1, EE19 USP20, EE STK11IP, and EE COG1) are shown in their genomic context, each predicted to contact its host or distal target genes via one or more interaction datasets (promoter capture Hi-C, ENCODE-rE2G, and GTEx). Arcs denote these predicted regulatory interactions. Bar graphs compare RT–qPCR measurements of gene expression in no-target controls (blue bars) versus CRISPRi-mediated EE silencing (yellow bars). EEs validated by both luciferase assays and STARR-seq (EE3 GARRE1, EE19 USP20) exhibit reduced transcription of both their host genes and predicted targets (b,c). Similarly, silencing of top-ranked EEs from STARR-seq (EE STK11IP, EE COG1) diminishes expression of associated loci (d,e). Error bars represent mean ± s.e.m. (n = 3 biological replicates). Statistical significance was assessed using a one-tailed independent t-test. Asterisks indicate p-values: *P < 0.05, **P < 0.01, ***P < 0.001, ns = not significant.

### Evolutionary dynamics and selective pressure on Exonic Enhancers

To determine whether EEs represent ancient, well-established regulatory elements or arise sporadically in the genome, we assessed their conservation patterns across multiple mammalian lineages and beyond. Pairwise genome alignments between human and mouse revealed that 28% of human-defined EEs map to mouse-defined EEs (Fig. 7a), indicating that a notable subset is evolutionarily retained. The remaining 70% appear human-specific, potentially reflecting lineage-specific regulation or incomplete annotation arising from differences in TF coverage across species. Notably, *shadow* EEs—exons in mouse that align to human-specific EEs—still display TF occupancy exceeding negative controls, suggesting emergent or sub-threshold enhancer potential. These *shadow* EEs could represent ancestral exonic enhancers that have partially decayed or remain undetected in some lineages, reflecting incomplete TF coverage or lineage-specific expansions. For comparison, we examined classic enhancers (Ctrl+), of which only 11% overlapped conserved enhancer loci in mouse, consistent with partial evolutionary retention of enhancer elements^23^ (Supplementary Fig. 18). Collectively, these findings suggest that EEs can follow both lineage-specific and conserved regulatory trajectories within coding sequences.

**Figure 7.**
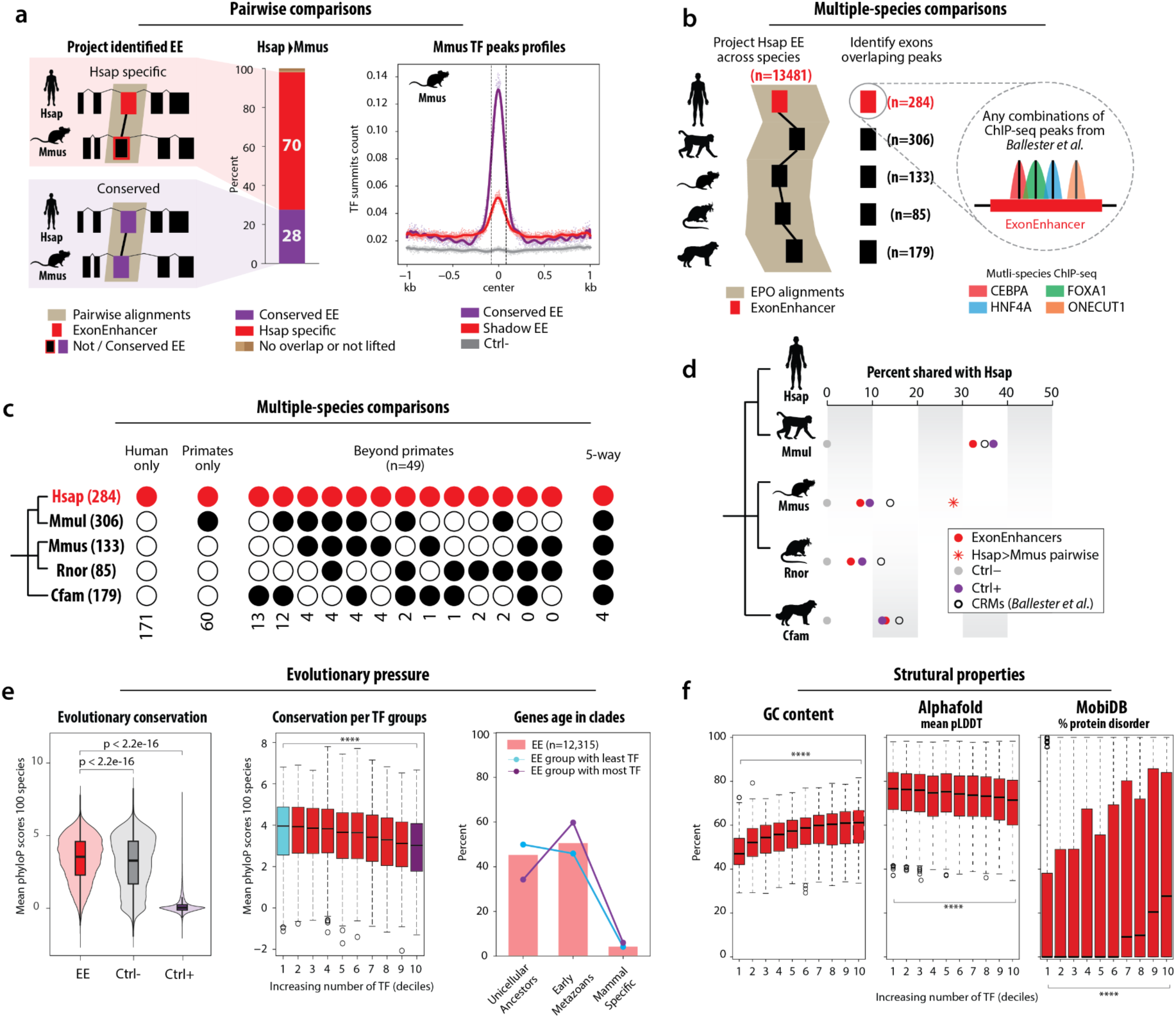
Evolutionary conservation and selective pressure on exonic enhancers across mammals and beyond. (**a**) Pairwise comparison of exonic enhancers (EEs) between human (*H. sapiens*, Hsap) and mouse (*M. musculus*, Mmus) using Ensembl pairwise genome alignments. Hsap-specific EEs (red) are those not conserved in the mouse genome, while conserved EEs (purple) meet the TF-binding criteria in both species. The bar plots quantify the fraction of Hsap EEs that either map to a corresponding Mmus EE or do not, and the TF summit profiles for *shadow* EEs in Mmus (red line) indicate potential enhancer-like features even in cases lacking formal EE annotation. (**b**) Schematic of a multi-species analysis leveraging data from Ballester et al., where four TFs (CEBPA, FOXA1, HNF4A, and ONECUT1) were profiled by ChIP-seq in five mammals (human, macaque, mouse, rat, and dog). Exonic regions exhibiting TF binding in each species were identified through multiple-genome alignments (EPO), permitting direct comparisons to Hsap EEs. (**c**) Multi-species EE occupancy analysis. Filled circles denote the presence of TF binding in that species. Black circles represent at least one TF peak, while red circles highlight the set of 113 human EEs that retain exonic enhancer features across multiple species. (**d**) Proportion of human exonic enhancers (EEs) shared with macaque (Mmul), mouse (Mmus), rat (Rnor), and dog (Cfam). Red circles indicate the percentage of EEs validated across multiple species analyses, while the red asterisk highlights the subset of EEs specifically conserved between human and mouse (panel a). Purple, grey, and open circles represent classic enhancers (Ctrl+), negative controls (Ctrl–), and cross-species conserved regulatory modules (CRMs) from Ballester *et al.*, respectively. (**e**) Evolutionary constraints on EE sequences. Box plots (left) show mean phyloP scores across 100-species alignments, revealing stronger conservation for EEs than for classic enhancers (Ctrl+). Further subdivision of EEs by increasing TF occupancy (middle) indicates that lowly bound EEs exhibit higher sequence constraint, while EEs with high TF occupancy display relatively lower constraint. Ancestral gene age analysis (right) suggests that low-TF EEs are more frequently associated with ancient gene families (unicellular ancestors), whereas high-TF EEs are often linked to later-evolving metazoan genes. (**f**) Structural properties of EEs grouped by TF occupancy (deciles, x-axis). Left: GC content (percentage) increases with higher TF occupancy (****P < 0.00005, Kruskal-Wallis test). Middle: Mean pLDDT scores from AlphaFold predictions decrease with increasing TF occupancy (****P < 0.00005, Kruskal-Wallis test), indicating lower confidence in protein structure prediction. Right: Percentage of EEs overlapping with predicted disordered protein regions (MobiDB) increases with TF occupancy (****P < 0.00005, Kruskal-Wallis test). Box plots show median (middle line), first and third quartiles (box), and 1.5× interquartile range (whiskers); outliers shown as individual points.

To extend these observations, we incorporated multi-species ChIP-seq data^24^ of four key TFs (CEBPA, FOXA1, HNF4A, and ONECUT1) in liver across human, macaque, mouse, rat, and dog. By intersecting EEs with these multi-genome TF ChIP-seq datasets and projecting them onto multiple alignments, we identified coding exons harbouring at least one TF peak in each species (Fig. 7b). Comprehensive comparisons (Fig. 7c) revealed consistent, though not ubiquitous, TF occupancy within exonic sequences across lineages, with some EEs spanning three or more species.

We next quantified the proportion of human EEs overlapping putative exonic enhancer sites in macaque, mouse, rat, and dog (Fig. 7d). This analysis confirmed that a subset of EEs (red circles) is shared across multiple species, whereas others remain lineage-specific. Notably, the fraction of EEs recapitulated in mouse (asterisk, comparing panel a) exceeded that observed in rat or dog, reflecting a partial correlation with phylogenetic distance. Comparisons to classic enhancers (Ctrl+) and cross-species conserved regulatory modules (CRMs^24^) revealed that EEs exhibit a similar pattern of partial conservation.

We next examined the nucleotide-level evolutionary conservation of EEs, both globally and stratified by TF occupancy (Fig. 7e). PhyloP scores from 100-species alignments^25^ revealed that EEs display significantly higher sequence constraint than control coding exons (Ctrl−) or typical enhancers (Ctrl+) (P < 2.2 × 10^−16, Wilcoxon rank-sum test). Moreover, EEs bound by fewer TFs tend to be more strongly conserved and associate with evolutionarily ancient genes with less variants, whereas those with higher TF occupancy exhibit signs of accelerated evolution, have more variants and associate with more recently evolved metazoan genes (Fig. 7e, middle and right, Supplementary Fig. 19).

To investigate the structural and biochemical properties of EEs, we analysed how GC content, 3D protein structure stability, and intrinsic disorder change with increasing TF occupancy in the four species (Fig. 7f, Supplementary Fig. 20). Grouping EEs by deciles of TF binding density revealed a positive correlation between TF occupancy and GC content, mirroring the reduced sequence conservation in highly bound EEs (Fig. 7e). Meanwhile, mean pLDDT scores (AlphaFold’s confidence metric for predicted 3D structure^26^) decreased with increasing TF occupancy, suggesting that highly bound EEs occur in structurally more flexible regions, potentially accommodating more TF-binding motifs or experiencing reduced structural constraints. MobiDB^27^ analysis further supported this notion, showing that highly TF-bound EEs are frequently located in intrinsically disordered protein segments. Overall, these data indicate that the molecular architecture of EEs balances transcription factor binding requirements with protein structural constraints.

## Discussion

Exonic enhancers (EEs) are emerging as dual-function elements that integrate within coding sequences to modulate transcription, challenging the traditional dichotomy between coding and noncoding genomic regions. In this study, we uncovered a widespread occurrence of EEs across multiple species, using large-scale TF-binding maps, open chromatin profiles, and enhancer-reporter assays. Our results indicate that, while many exons merely encode protein domains, a substantial subset also harbours hallmark enhancer features—DNase hypersensitivity, histone modifications, G-quadruplex formation, and TF co-occupancy—underscoring their capacity to regulate gene expression.

Beyond their prevalence, EEs influence key biological processes. Functional assays, including STARR-seq and luciferase reporter assays, confirmed that EEs drive transcription in cis. Notably, these assays measured enhancer activity solely from the exonic sequences, demonstrating that the exon itself, and not its intronic flanking regions, harbours cis-regulatory potential. Furthermore, promoter capture Hi-C and ENCODE-rE2G data reveal that many EEs physically interact with gene promoters, consistent with classical enhancer looping. In agreement, CRISPRi experiments targeting selected EEs confirmed their regulatory function on both host and distal neighbouring genes. Mechanistically, these EE elements may integrate multiple TF inputs within coding regions, effectively bridging the coding and regulatory layers to coordinate gene expression.

Our investigation also highlights the potential for EEs to shape disease phenotypes. By intersecting EEs with genomic variation datasets (e.g., gnomAD, TCGA), we identified both common and rare variants that overlap critical TF-binding sites. These mutations, ranging from synonymous to nonsynonymous, can markedly alter EE-driven enhancer activity in reporter assays, suggesting that even “silent” coding changes may have regulatory consequences. Moreover, exons with weak or no enhancer activity could acquire regulatory functions through the incorporation of novel variants. In cancer, EE variants were associated with differential expression of putative target genes, which in turn correlated with patient survival. These findings expand emerging evidence that coding-region mutations can exert non-canonical effects on gene regulation, thereby influencing cancer progression and other disease processes.

From an evolutionary perspective, EEs display a complex pattern of conservation and innovation across vertebrate lineages. Approximately 28% of human EEs show conserved regulatory signatures in mouse, while others exhibit lineage-specific patterns, suggesting dynamic regulatory evolution. The elevated sequence constraint observed in EEs, compared to typical coding exons or classical enhancers, indicates strong purifying selection to maintain both coding and regulatory functions. Moreover, the correlation between TF occupancy and gene age suggests distinct evolutionary trajectories: ancient genes tend to harbour EEs with lower TF complexity, whereas metazoan-specific genes often contain EEs bound by multiple factors. It remains an open question whether coding exons acquire enhancer-like activity through variant acquisition, or whether pre-existing intronic enhancers accumulate mutations that eventually incorporate them into coding sequences.

These findings necessitate a fundamental change in interpreting genetic variation in coding regions. The dual functionality of EEs implies that coding-sequence mutations should be evaluated not only for their impact on protein structure but also for potential regulatory consequences. This perspective has immediate implications for variant-interpretation pipelines, which must extend to coding exons to detect potential enhancer disruptions in research and clinical settings, particularly in cancer genomics where coding mutations are routinely assessed for pathogenicity.

Looking ahead, investigating the tissue-specific or cell type–specific contexts in which EEs operate will be critical to understanding their broader biological impact. Future efforts should also extend cross-species comparisons to reveal the evolutionary forces shaping EEs over longer timescales. By highlighting their prevalence, functionality, and pathological significance, our findings underscore a conceptual shift: exons are not merely the blueprint for proteins, but also an integral part of the regulatory machinery driving precise spatiotemporal gene expression.

## Methods

### Genomic annotation resources

Annotations for coding exons and transcripts were obtained from the UCSC Table Browser^25^ using GENCODE^28^ v41 (hg38) for human (*Homo sapiens*), GENCODE M33 (mm39) for mouse (*Mus musculus*), Ensembl^29^ Release 104 (BDGP6/dm6) for fruit fly (*Drosophila melanogaster*), and Ensembl Release 44 (TAIR10) for *Arabidopsis thaliana*. For Transcription Start Site (TSS) data, FANTOM5^9^ TSS peaks were used for human and mouse, with mm10 TSS peaks converted to mm39 using UCSC LiftOver. For Drosophila melanogaster, TSS peaks were obtained from ModENCODE^30^ CAGE-seq experiments (NCBI accession number SRP001602) using the GEP UCSC Table Browser. For *Arabidopsis thaliana*, a comprehensive TAIR10 TSS catalogue was generated by integrating results from these studies^31–33^. Transcriptional regulators binding catalogues were obtained from ReMap^5^ (https://remap.univ-amu.fr/, version 2022), which provides a curated and integrated catalogue of TF ChIP-seq data across multiple species.

### Defining and characterizing Exonic Enhancers

To systematically identify exonic enhancers (EEs), we first generated a comprehensive set of protein-coding exons using GENCODE v41 (human), GENCODE vM33 (mouse), FlyBase r6.54 (D. melanogaster), and Araport11 (A. thaliana) annotations. Overlapping isoform exons with a size difference of ≤50 bp were merged to avoid fragmenting potential regulatory elements. To minimize confounding signals from promoter-proximal regions, we removed exons within 1 kb (human, mouse) or 100 bp (fly, Arabidopsis) of annotated transcription start sites (TSS) or termination sites (TES) in the same orientation. We further filtered out exons overlapping TSS peaks within 500 bp (human, mouse) or 100 bp (fly, Arabidopsis) windows. EE candidates were defined using transcription factor ChIP-seq peaks from ReMap. Candidate selection required: (i) a minimum of 10 distinct TFs overlapping the exon, (ii) the maximum TF density within the coding exon must exceed the local TF density measured in the ±50 bp flanking exonic region, to avoid artificially capturing TF clusters near exon–intron boundaries, and (iii) a minimum ratio of 0.5 between TF peak summits and total TF peaks overlapping the exon (Supplementary Fig. 21). For human, additional filtering was applied to refine EE candidates. Only exons from transcripts with a transcript support level (TSL) of 1 were retained, ensuring high-confidence annotations.

Furthermore, for human and mouse, exons overlapping promoter-associated chromatin marks from ENCODE cCREs were removed. In the case of the mouse genome, mm10 promoter marks were lifted to mm39 using UCSC LiftOver to maintain consistency across genome builds.

For control datasets, negative controls were defined as protein-coding exons that passed the same filtering process as EEs but had no overlapping TFs. Positive controls consisted of intergenic distal enhancers from ENCODE cCREs^6^ (human, mouse) or the literature (fly, Arabidopsis) that passed the same filtering process as EEs and overlapped at least 10 TFs, ensuring that control elements were subject to equivalent selection criteria.

### DNase-seq and ATAC-seq data processing

We compiled a comprehensive atlas of chromatin accessibility data from multiple sources. For human (GRCh38), DNase-seq peaks were obtained from ENCODE through the UCSC Genome Browser database. Mouse DNase-seq data (ENCODE accession ENCFF910SRW) was lifted from mm10 to mm39 using UCSC liftOver (minMatch=0.95). For both human and mouse, as well as D. melanogaster (dm6), we retrieved DNase-seq and ATAC-seq peaks with a significance threshold of 50 from ChIP-Atlas^8^. For A. thaliana (TAIR10), DNase-seq data was obtained from PlantRegMap^34^. ATAC-seq data was collected from the Gene Expression Omnibus under accessions GSE101482, GSE101940, GSE122772, GSE164159, and GSE173834. Processed datasets for the four species are provided in a Supplementary file Omics. All peak coordinates were standardized to their respective reference genome builds: GRCh38 (human), GRCm39 (mouse), dm6 (D. melanogaster), and TAIR10 (A. thaliana).

### STARR-seq atlas processing and analysis

We constructed a comprehensive STARR-seq atlas by curating 112 published datasets, and by compiling active (log2FC > 0) and inactive (log2FC ≤ 0) peaks across four species (H. sapiens, M. musculus, D. melanogaster, and A. thaliana). All STARR-seq peaks were standardized to ENCODE narrowPeak format. Peak coordinates from earlier genome builds were converted to current assemblies (GRCh38, GRCm39, dm6, TAIR10) using UCSC liftOver (minMatch=0.95). Statistical significance was assessed using reported q-values or p-values where available (P < 0.05). A complete list of datasets and their sources is provided in Supplementary file STARR-seq catalogues. The biotype signatures of exonic enhancers (EEs) were determined by first assigning each ChIP-seq cell line to a biotype as previously described^35^, followed by calculating the normalized abundance of each biotype across EEs. For K-562 and A-549 STARR-seq analysis, we generated a composite signal track using deepTools^36^ v3.5.1. Specifically, fold-change signals from replicates were computed using bigwigCompare, then merged using bigWigMerge. Used datasets for the signal tracks are also listed in the Supplementary file STARR-seq catalogues.

### G-quadruplex analysis

Potential G-quadruplex (G4) forming sequences were identified using G4-seq data generated by an optimized G4 sequencing method^37^. The G4 peak data were retrieved from the Gene Expression Omnibus (GEO) under accession number GSE110582. The enrichment of G4 structures in EEs compared to control regions (Ctrl+ and Ctrl-) was assessed using Fisher’s exact test (one-sided, P < 0.05).

### TF peaks and TF binding site matching and randomization

For each EE, we quantified the overlap (“correspondence rate”) between transcription factor (TF) fragments (score ≥ median) intersecting the EE and their corresponding transcription factor binding sites (TFBSs) derived from the JASPAR2022 database^19^ (https://jaspar.elixir.no/). These TF peaks were then randomly redistributed across the genome to generate a null distribution of correspondence rates. The observed correspondence rate in EEs was subsequently compared with the rate obtained under randomization.

### Selection of candidate EEs for luciferase assays

Candidate EEs for luciferase assays were selected based on the presence of K-562 STARR-seq and DNase-seq peaks. Single-nucleotide polymorphism (SNP) data in the hg38 reference genome were obtained from gnomAD^18^ v3 (https://gnomad.broadinstitute.org/), and their potential impact on transcription factor (TF) binding was predicted using the FABIAN-variant tool^38^. Where necessary, synonymous SNPs were manually introduced for construct generation. Missense and synonymous variants tested for enhancer activity gain or loss were chosen based on their predicted effect on existing TFs and TFBSs within each candidate EE.

### Cell culture

The K-562 cell line (ATCC CCL-243), derived from chronic myelogenous leukemia, was obtained from the American Type Culture Collection (ATCC) and cultured in RPMI 1640 medium with GlutaMAX (Thermo Fisher Scientific), supplemented with 10% fetal bovine serum (FBS; Thermo Fisher Scientific). Cells were incubated at 37 °C in a humidified atmosphere containing 5% CO₂, passaged every three days at a density of 2 × 10⁵ cells/mL, and routinely tested for mycoplasma contamination.

### Experimental Validation of Exon Enhancer Activity

We tested 23 sequences corresponding to identified exonic enhancers (Supplementary Table S1-3). Each sequence (lucEE1 to lucEE23) was cloned into the pGL3-Promoter vector (Promega, E1761), containing the SV40 promoter upstream of the luciferase gene, at the BamHI restriction site. For transfection, 1 × 10⁶ K-562 cells were electroporated with 1 µg of each lucEE construct and 200 ng of Renilla control vector using the Neon Transfection System (Thermo Fisher Scientific; pulse voltage: 1450 V, pulse width: 10 ms, pulse number: 3) and cultured in 1 mL of medium in 24-well plates. After 24 h, cells were lysed, and luciferase activity was measured using the Dual-Luciferase Reporter Assay (Promega). Firefly luciferase data were normalized to Renilla luciferase activity, and results were expressed as fold-change in relative light units compared to the empty pGL3-Promoter vector. Experiments were performed in triplicate and repeated three times.

### Exon Enhancer Activity Gain or Loss Assays

To assess the impact of sequence modifications on enhancer activity, the same protocol described above was applied, using synthetic exon DNA fragments containing designed sequence-specific modifications (Supplementary Table S2). Luciferase activity was measured to determine whether these alterations led to gain or loss of function in enhancer activity.

### Luciferase constructs for Promoter–Enhancer evaluation

To test promoter activity, DNA fragments corresponding to the promoter regions of the GARRE1 gene and its partner genes (GPI and PDCD2L), or the USP20 gene and its associated partners (C9orf78 and TOR1B), were synthesized and cloned into the MluI and XhoI sites upstream of the luciferase coding sequence, generating six reporter constructs (Supplementary Table S2). For each promoter construct, the corresponding wild type exonic enhancer (EE3_WT or EE19_WT) was cloned into the BamHI restriction site (Supplementary Table S3). Luciferase assays were performed following the same protocol described above for exonic enhancer activity assays.

### EE STARR-seq processing and analysis

#### Selection Criteria

EE tested in the STARR-seq experiments were primarily identified based on signatures derived from the K-562 cell line, complemented by additional EEs exhibiting signatures in GM12878 and A-549 cells. Selection criteria included: (i) at least one non-redundant TF peak summit overlapping a DNase-seq peak, and (ii) at least one non-redundant TF peak summit coinciding with at least three enhancer peaks identified in the STARR-seq catalogue. Only EEs ≥130 bp were retained. For EEs exceeding 220 bp, the sequence was trimmed to a 220 bp region centered on the highest concentration of TF peaks, remaining within the exon boundaries (Supplementary Fig. 22a).

#### SNP-Modified Sequences

For K-562, single-SNP mutant constructs were generated by introducing synonymous variants selected from gnomAD v3 (https://gnomad.broadinstitute.org/) that were predicted to alter TF/TFBS binding by more than |0.66|, as estimated by the FABIAN-variant tool. Multi-SNP mutant constructs were generated in the same manner, but included at least two additional non-adjacent SNPs per sequence.

#### Controls

Negative controls were defined as regions lacking any ATAC-seq or DNase-seq peaks and exhibiting the lowest JASPAR2022 TFBS scores. Positive controls were randomly drawn from regions containing at least three TF peaks in each of the three cell lines. All control fragments were 130–220 bp in length. Additionally, 200 negative and positive “STARR+/-” technical controls were included based on published STARR-seq experiments^20^. The complete list of all 11,999 tested sequences is provided in Supplementary file STARR-seq.

### STARR-seq Library Construction and Transfection

A total of 11,999 candidate sequences were synthesized by Twist Bioscience, each appended with 5′-ACGCTCTTCCGATCT and 3′-AGATCGGAAGAGCAC, which are 15 bp invariable adapters corresponding to TruSeq read indices. The resulting oligo library was resuspended in H₂O at 10 ng/µL. To add homology arms for subsequent cloning (PCR_A), 10 ng of the oligo library was amplified in three parallel 50 µL reactions using KAPA HiFi Hotstart Readymix (Roche, cat. #07958927001), 10 µM forward primer Fw_Enh_MPRA (CAACTGATCTAGAGCATGCAACGCTCTTCCGATCT), 10 µM reverse primer Rv_Enh_MPRA (GAAGCGGCCGGCCGAATTCGTGTGCTCTTCCGATCT), and the following program: 98 °C for 2 min; 20 cycles of (98 °C for 20 s, 60 °C for 30 s, 72 °C for 30 s); and a final extension at 72 °C for 2 min. The three PCR products were purified with Nucleospin cleanup (Macherey-Nagel, cat. #740609.50) in a 20 µL elution, then desalted on a MF-Millipore MCE 0.025 µm membrane (Sigma, cat. #VSWP02500) for 25 min.

The STARR-seq vector was digested with AgeI-HF, SalI-HF, and NotI-HF (NEB, cat. #R3552S, #R3138S, #R3189S), loaded onto a 1.5% agarose gel at 100 V for 45 min, and the desired band was extracted using the Nucleospin Gel Cleanup kit (Macherey-Nagel, cat. #740609.50). For Gibson Assembly (NEB, cat. #E5510S), 150 ng of digested vector and 40 ng of the purified PCR_A product were incubated in two separate 20 µL reactions at 50 °C for 45 min. The assembled products were then desalted on a MF-Millipore MCE 0.025 µm membrane for 25 min.

#### Transformation and Library Expansion

Each 15 µL Gibson assembly reaction was electroporated into 40 µL of MegaX DH10B T1R cells (Thermo Fisher, cat. #C640003) using a Bio-Rad Gene Pulser II system (2 kV, 200 Ω, 25 µF) in 0.1 cm cuvettes. Cells were recovered for 1 h at 37 °C in 8 mL SOC medium. To estimate library coverage, 1 µL of the recovery culture was plated on LB + carbenicillin (Invitrogen, cat. #10177-012) and grown overnight at 37 °C, yielding ∼775 colonies per µL (516× coverage for 12,000 inserts). The remaining transformation was spread over eight 15 cm LB + carbenicillin plates and grown for 16 h at 37 °C. Colonies were then transferred to 1 L of Terrific Broth with 1× carbenicillin and incubated at 37 °C, 100 rpm for 16 h. Ten midipreps (Nucleobond Midiprep Kit, Macherey-Nagel, cat. #740410) were combined, eluted in a total of 500 µL, and yielded 1,751 µg of Exonhancer plasmid library DNA.

#### Cell Culture and Transfections

K-562 cells were maintained in RPMI 1640 (Thermo Fisher) and passed for at least one week prior to transfection, with 97% viability at the time of use (passage +10). For transfection, 50 × 10⁶ cells were pelleted, washed in PBS, and resuspended in 1 mL Neon resuspension buffer (Thermo Fisher). A total of 250 µg Exonhancer plasmid library was added, and 10 transfections (100 µL each) were performed (settings: 1450 V, 10 ms, 3 pulses). Two transfections (200 µL total) were combined per replicate in a culture dish with 10 mL of RPMI 1640 and incubated at 37 °C for 24 h, creating five replicates. As a control, one replicate was transfected using the STARR-seq empty vector using the same settings. 24 hours after transfection, GFP was assessed as an indication of enhancer activity (Supplementary Fig. 22c).

#### gDNA Isolation and STARR-seq Protocol

After 24 h of incubation, gDNA was extracted from 1 mL of each of three replicates using the Flexigene protocol in a final volume of 50 µL to assess input library integrity. These gDNA samples, plus a midiprep reference, were each PCR-amplified (10 ng input) using KAPA HiFi (50 µL reactions) with primers Fw #11 (GGGCCAGCTGTTGGGGTGAGTAC) and Rv #10 (CTTATCATGTCTGCTCGAAGC) under the following program: 98 °C for 2 min; 15 cycles of (98 °C for 20 s, 65 °C for 20 s, 72 °C for 1 min); 72 °C for 2 min. Products (∼1350 bp) were purified (Nucleospin) and quantified by dsDNA HS Qubit.

For the STARR-seq assay, total RNA was extracted (RNeasy, Qiagen), followed by mRNA isolation (Dynabeads, Invitrogen) for each of the 5 replicates. cDNA was synthesized using Superscript III (Thermo Fisher) with a reverse transcription primer (CAACTCATCAATGTATCTTATCATG) and an RNase H step at 37 °C for 20 min. cDNA was purified, quantified via ssDNA Qubit, and amplified (PCR1) using KAPA HiFi and primers Fw (GGGCCAGCTGTTGGGGTGTCCAC) and Rv (CTTATCATGTCTGCTCGAAGC) under the program: 98 °C for 2 min; 15 cycles of (98 °C for 20 s, 65 °C for 20 s, 72 °C for 1 min); and 72 °C for 2 min. A second indexing PCR (dual-index TruSeq) was conducted (20 cycles) on PCR1 products from both gDNA and cDNA samples, followed by purification (AmpureXP) and verification of product size (240–360 bp) by Qubit and Bioanalyzer. Libraries were sequenced on a NextSeq 2000 (Illumina).

### STARR-seq processing

Sequencing paired-end reads were aligned to the reference library using Bowtie2 with the “very-sensitive” parameter set to ensure high-fidelity mapping. Resulting BAM files were processed with SAMtools^39^ to remove unmapped reads and any reads aligned to the reverse strand (using samtools view -F 20). The filtered reads were then sorted and indexed, and unique identifiers in the library were quantified to produce raw read counts for each sample. Counts were normalized to counts per million (CPM) to enable cross-sample comparison. Finally, the pipeline computed the ratio and log₂ ratio of normalized cDNA counts to the corresponding input sample. Sequences with an input read count < 500 or with an input STD >= 20 were discarded (Supplementary Fig. 22b). SNP-associated effects on enhancer activity in K-562 cells were assessed according to Long *et al.* method^40^ using a two-sided Wald test, with significance set at FDR < 0.05 (Benjamini–Hochberg correction).

### CRISPRi targeting of EEs

#### Generation of CRISPRi-competent K-562 cells

A CRISPRi-competent K-562 clone was generated by transducing cells with a dCas9–KRAB–MeCP2 lentiviral vector (Addgene #122205). Lentiviruses were produced in HEK293T cells co-transfected with the VSVG packaging plasmid via calcium phosphate precipitation. K-562 cells (0.3 × 10⁶ cells/mL) were infected twice in the presence of blasticidin selection. Single cells were seeded into 96-well plates were subsequently infected with a Crop-seq-derived lentivirus (Addgene #86708) encoding an sgRNA targeting the human CD81 promoter (Supplementary Table S4), followed by seven days of puromycin selection. Individual clones were assessed for CD81 surface expression by cytometrie and validated by RT-qPCR. A clone exhibiting efficient CD81 knockdown was chosen for further CRISPRi experiments (Supplementary Fig. S23).

#### CRISPRi targeting of EEs

sgRNAs against selected EEs were designed using the CRISPOR tool^41^, synthesized, and cloned into the Crop-seq-guide-puro vector (Genecust, Luxembourg) via the BsmBI site. A no-target sgRNA was also cloned as a control. Lentiviral vectors were amplified in Endura E. coli cells (Lucigen), and lentiviral particles were produced as described above. For each EE, 2 × 10⁴ K-562 cells expressing dCas9–KRAB–MeCP2 were independently transduced at a multiplicity of infection (MOI) of 10 with 2 mL of complete RPMI medium supplemented with 10% FBS and 1× Penicillin– Streptomycin–Glutamine (Thermo Fisher). Ten days post-infection, cells were harvested for RNA extraction. The list of sgRNA is provided in Supplementary Table S4.

#### Gene expression analysis

Total RNA was extracted using the RNeasy Plus Mini Kit (Qiagen) following the manufacturer’s protocol. One microgram of RNA was reverse-transcribed with SuperScript™ VILO™ Master Mix (Thermo Fisher Scientific, #11755250). qRT–PCR was performed using SYBR Green Master Mix (Thermo Fisher Scientific) on a QuantStudio™ 6 Flex (Thermo Fisher Scientific) instrument, with 1:10 diluted cDNA. Relative expression was analysed by the 2^ΔΔCT method, normalized to GAPDH expression. Each group was tested in three independent RNA and cDNA preparations, and the mean ± standard deviation was calculated relative to the no-target control. Primers used are listed in Supplementary Table S5.

### Integration of Hi-C, ENCODE-rE2G, and eQTL data for EE interaction analysis

To identify putative enhancer–promoter interactions, we overlapped EEs mapped to the hg38 reference genome with the promoter capture Hi-C dataset^16^. We retained cases in which (i) an EE resided in a target fragment interacting with a complementary fragment (n=9340), or (ii) an EE lay within a baited (promoter) fragment interacting with another baited fragment (n=1637). EEs were further overlapped with predicted regulatory interactions from the ENCODE-rE2G^17^ model using bedtools^42^ intersect (option *-f 0.50*). Additionally, we intersected EEs with expression quantitative trait loci (eQTLs) from GTEx^43^ v8. We defined an “internal interaction” as an EE that targets a gene located within its own gene body, whereas “external interactions” refer to EEs targeting genes outside their host gene body. Finally, “robust interactions” were those for which at least two of the above datasets (promoter capture Hi-C, ENCODE-rE2G, eQTLs) provided support.

### Impact of Exonic Enhancer mutations in cancer: TCGA PanCanAtlas Analysis

#### Mutation Data Processing

Somatic mutation data from The Cancer Genome Atlas (TCGA) were retrieved from the publicly available multi-cancer mutation annotation format (MAF, [mc3 v.0.2.8](https://gdc.cancer.gov/about-data/publications/mc3-2017)) file provided by the TCGA PanCanAtlas group^44^. To integrate these data with EEs, genomic coordinates of EEs were lifted over from hg38 to hg19 using UCSC LiftOver, followed by bedtools intersect to identify overlaps between EEs and somatic mutations.

#### Differential Expression Analysis of EE-Target Genes

To assess the impact of EE mutations on gene expression, we analysed differential expression in genes with robust EE–target interactions (defined as interactions supported by at least two datasets: promoter capture Hi-C, ENCODE-rE2G, or GTEx eQTLs). Differential expression analysis was performed using pyDESeq2^45^ (v0.4.8), comparing two groups of patients: i) Mutant Group: Patients carrying mutations within a given EE, ii) Wild-type Group: Patients with no mutations in the corresponding EE.

A separate analysis was conducted where only patients carrying synonymous (silent) mutations within a given EE were retained and compared against those without mutations. Differentially expressed genes were identified based on an absolute fold-change threshold of |FC| > 0.5 and a significance cutoff of Padj < 0.05 (Benjamini–Hochberg correction). All EE mutation annotations and associated expression changes are provided in the Zenodo repository (https://doi.org/10.5281/zenodo.15079251) and in the Supplementary table.

### PanCan survival analyses

To assess the potential clinical impact of EE-mediated gene dysregulation, we selected differentially expressed EE-target genes identified in four cancer types: lower-grade glioma (LGG), lung adenocarcinoma (LUAD), stomach adenocarcinoma (STAD), and skin cutaneous melanoma (SKCM). These genes were chosen based on their differential expression patterns in TCGA PanCanAtlas data (see "TCGA PanCanAtlas Analysis" section) and their classification as robust EE targets (i.e., supported by at least two of the following datasets: promoter capture Hi-C, ENCODE-rE2G, or GTEx eQTLs). To evaluate the prognostic significance of EE-target gene expression, we used the cSurvival^46^ tool (v1.0.6). Kaplan–Meier survival curves were generated for high- and low-expression groups, stratified based on the median expression of each target gene across cancer patients. Log-rank tests were applied to determine statistical significance (*P* < 0.05). For each cancer type, we assessed the overall survival (OS) of patients grouped according to target gene expression levels. Hazard ratios (HRs) and confidence intervals (CIs) were computed using a Cox proportional hazards model, adjusting for relevant clinical covariates such as tumour stage and patient age when available.

### Variant effects predictions and visualization in Exonic Enhancers

To assess how single-nucleotide polymorphisms (SNPs) may alter transcription factor (TF) binding within exonic enhancers (EEs), we used the FABIAN-variant^38^ tool. FABIAN-variant quantifies binding affinity changes by evaluating the presence or absence of TF motifs at SNP-affected sites. For each variant (TCGA, gnomAD, Supplementary Fig. 24) located within an EE, we computed its predicted effect on associated TFBSs. Mutations that significantly altered TF-binding affinity (binding gain or loss) were recorded. To facilitate the interpretation of TF-binding alterations, we visualized the computed TF-binding impacts using lollipop plots. These plots were generated using track visualization tools in UCSC Genome Browser^25^. These visualizations provide a systematic overview of how mutations in exonic enhancers may alter TF-binding landscapes, with potential implications for gene regulation in cancer. The full lollipop track illustrating these TFBS alterations is available in a public trackhub and sessions listed at UCSC (sessions: https://genome.ucsc.edu/s/Benoit%20Ballester/hg38_ExonEnhancers_gnomAD; https://genome.ucsc.edu/s/Benoit%20Ballester/hg19_ExonEnhancers_TCGA).

### Processing GWAS Catalog and 1000 Genomes SNP Data

GWAS variants from the NHGRI-EBI GWAS Catalog^47^ (v1.0.2) and common human SNPs (in VCF format) from the 1000 Genomes Project^48^ were retrieved. GWAS variants without rsIDs or genomic coordinates were removed. Common SNPs were filtered in PLINK (v1.9) using the parameters --maf 0.01, --geno 0.05, and --hwe 1e-6. Next, common SNPs within 1 Mb of each GWAS lead SNP and showing high linkage disequilibrium (r² > 0.8) were identified with PLINK (–ld-window-kb 1000 –ld-window-r2 0.8). Each GWAS trait was then mapped to one of the parent trait categories—“Disease,” “Biological process,” “Measurement,” “Phenotype,” or “Other”—based on the Experimental Factor Ontology (EFO). Finally, we normalized the number of SNPs in each category by the total number of elements in the overlapped dataset (EE or controls).

### Conservation and Evolutionary Comparisons

Cross-Species Projection of Exonic Enhancers (EEs) was performed using both pairwise and multiple genome alignments. For the pairwise approach, EEs defined in the human genome (hg38) were mapped to the mouse genome (mm39) through the Ensembl API with LASTZ-net alignments (Ensembl release 104). High-confidence mappings were obtained by retaining regions with at least 90% sequence identity over 50 bp, and coordinate conversion was conducted using a chain-based liftOver. For the multiple genome alignment approach, EEs in hg38 were projected onto mm39 (Mus musculus), mml10 (Macaca mulatta), rnor7 (Rattus norvegicus), and cfam1 (Canis lupus familiaris) via the Enredo-Pecan-Ortheus (EPO^49^) alignments. Coordinate conversion blocks were retrieved from Ensembl, and in-house Perl scripts (v3.8) were used to identify orthologous exonic regions in each genome.

### Transcription Factor ChIP-seq Analysis Across Species

ChIP-seq data was retrieved from Ballester et al.^24^, which profiled four key transcription factors (CEBPA, FOXA1, HNF4A, and ONECUT1) in the liver of human, macaque, mouse, rat, and dog. Reads were pooled prior to alignment and were trimmed and filtered using fastp (v.0.23.1) with the “--max_len1 36 --cut_tail --cut_tail_mean_quality 25 --average_qual 20” parameters. Reads were aligned to the respective genomes and outputted in BAM format using bowtie2 (v.2.5.1) and samtools view (v.1.18) under default parameters. Unmapped, duplicate, and multimapping reads were removed using sambamba (v.1.0.0). Peak calling was performed with MACS2 (v.2.2.7.1) callpeak, applying the effective genome sizes (“-g” parameter) computed with unique-kmers.py from the khmer^50^ program. The resulting TF peak sets were then intersected with the projected EEs using BEDTools (v.2.30.0), and cross-species enhancer occupancy was assessed based on regions that overlapped at least one TF peak in each lineage.

### Computing Evolutionary Conservation Scores

PhyloP conservation scores were obtained from multiple sources: human (hg38) 100-species phyloP scores, mouse (mm39) 35-species scores, and Drosophila melanogaster (dm6) 124-species scores were retrieved from the UCSC Genome Browser, while Arabidopsis thaliana (TAIR10) 63-species scores were acquired from PlantRegMap^34^. Mean phyloP scores for each EE were calculated using bigWigAverageOverBed (Kent utilities suite, v2), excluding regions that lacked phyloP coverage. Comparisons of EE conservation scores were made against control coding exons (without enhancer activity) and classic intergenic enhancers. The Kruskal–Wallis test (P < 0.05) was used to compare phyloP conservation among EEs, focusing on the first and last deciles of each distribution.

### Gene Age Analysis

Gene age categories were retrieved from Trigos et al.^51^ (reference). EEs were mapped to their host genes based on GENCODE (release 41) annotations, and each gene was classified according to its evolutionary origin. EEs were stratified by gene age category (e.g., unicellular, metazoan), and enrichment analyses were used to determine whether TF-bound EEs were distributed preferentially within specific evolutionary strata.

### EEs in Protein Structures

Amino acid sequences of exonic enhancers (EEs) in their corresponding host genes were retrieved for four species with geno2proteo^52^ R package (v.0.0.6). For each gene, the EE amino acid sequences were aligned to full-length protein sequences contained in AlphaFold^26^ CIF files and MobiDB^27^ entries, prioritizing the UniProtKB/Swiss-Prot identifier, then UniProtKB/TrEMBL, and finally the gene name. After mapping, we computed the average AlphaFold pLDDT score (reflecting predicted structural confidence) and the MobiDB-predicted mean disorder rate (reflecting structural flexibility) for each EE. Statistical significance of differences in these measures between the first and last deciles of EEs was assessed using the Kruskal–Wallis test (P < 0.05).

## Supporting information

Supplementary file omics

Supplementary file starrseq catalogues

Supplementary file starrseq experiment

Supplementary file tcga pancan

Supplementary file bed

## Data availability

Data supporting the findings of this study are available in a Zenodo repository https://doi.org/10.5281/zenodo.15079251, organized by analysis block in accordance with the structure of the paper. All datasets used in this study are publicly available or have been deposited in appropriate repositories. Annotated coding exons and transcripts (human, mouse, fly, thale cress) were retrieved from UCSC and Ensembl, while FANTOM5, modENCODE, and Arabidopsis TSS data are detailed in the Methods. ChIP-seq data for exonic enhancers (ReMap2022), DNase-seq and ATAC-seq data (ENCODE, ChIP-Atlas, PlantRegMap), and STARR-seq catalogue (Supplementary file STARR-seq catalogues) were integrated to identify EEs. G-quadruplex sequencing data (GSE110582) and the newly generated EE STARR-seq dataset (GEO accession GSE292804) were also incorporated. Cancer mutation data were obtained from the TCGA PanCanAtlas. Genomic interactions (promoter capture Hi-C), eQTLs (GTEx v8), and ENCODE-rE2G mappings were used to define EE–gene relationships, while phyloP conservation scores (UCSC, PlantRegMap) and gene age classifications (Trigos et al.) further contextualized EE evolution.

## Code availability

We deposited codes and bioinformatics environments in GitHub at https://github.com/benoitballester/ExonEnhancer. Processed data and files are available at Zenodo repository https://doi.org/10.5281/zenodo.15079251. Both the data and codes are publicly available for the replication of the whole study.

## Acknowledgements

The authors thank Robin Steinhaus for his assistance with fabian-tools and lifting the VCF query capacity. We also appreciate Science AAAS for granting permission to reproduce and modify the PanCanAtlas schema in Fig. 5c. This work was supported by a PhD Fellowship awarded to J.-C.M. from the French Ministry of Higher Education and Research (MESR), Institut National de la Santé et de la Recherche Médicale (INSERM), the Core Cluster of the Institut Français de Bioinformatique (IFB; ANR-11-INBS-0013), and by the Agence Nationale pour la Recherche (ANR; grant ANR-23-CE12-0008-01). A Marie Sklodowska-Curie Action postdoctoral fellowship (Eprom-101065610) supported A.V.O. We acknowledge the contribution of the SFR Biosciences (UAR3444/CNRS, US8/Inserm, ENS de Lyon, UCBL) facilities, particularly the AniRA lentivectors production facility of the CELPHEDIA Infrastructure (Gisèle Froment and Caroline Costa). We thank the Marseille-Luminy cell biology platform for managing cell culture and Nori Sadouni from HL BIOPROCESS (Marseille, France) for the STARR-seq preprocessing. The results presented here are based on data generated by the TCGA Research Network, the GTEx project, the ENCODE Consortium and its production laboratories, as well as independent laboratories that submitted raw ChIP-seq and other omics datasets to public repositories (GEO).

## Contributions

B.B. conceived and supervised the project. J-C.M. developed computational methods, curated ATAC-seq, DNase I, and STARR-seq datasets, and performed data analysis. M.T. carried out luciferase reporter assays. I.M.S. and M.T. conducted CRISPRi experiments. A.V.O. performed STARR-seq assays and selected and designed CRISPRi guides. S.S. supervised STARR-seq and CRISPRi experiments. J-C.M. and B.B. prepared the figures, and J-C.M., S.S., and B.B. wrote the manuscript with input from all authors.

