## Supplementary information for "Exonic enhancers are a widespread class of dual-function regulatory elements"

**Supplementary tables** 2
Table S1: EE tested by reporter luciferase assay Figure 3a. 2
Table S2: Tested promoters on Figure 3bc. 2
Table S3: EE mutated on Figure 3de. 3
Table S4: gRNAs used for CRISPRi Figure 6b-e. 3
Table S5: Primers to assess gene expression Figure 6b-e. 4

**Supplementary figures** 5
Figure S1: Re-evaluation of exon enhancer candidates from Birnbaum et al. 5
Figure S2: Genomic distribution of ChIP-seq and DNase-seq data across four species. 6
Figure S3: Transcription factor occupancy in coding exons across species. 7
Figure S4: Genomic distribution of transcription factor binding across exon classes. 8
Figure S5: Transcription factor binding density in internal exons. 9
Figure S6: Enrichment of transcription factors in EEs 10
Figure S7: Prevalence of exon enhancers across gene transcript isoforms. 11
Figure S8: Chromatin accessibility at exon enhancers across species. 12
Figure S9: Enrichment of histone modifications at exon enhancers. 13
Figure S10: STARR-seq activity in exonic enhancers stratified by biotype 14
Figure S11: Transcription factor binding site density at exon enhancers. 15
Figure S12: GC content and CpG enrichment at exon enhancers. 16
Figure S13: Cell line specificity and selection refinement of exon enhancer activity in STARR-seq. 17
Figure S14: Interaction landscape of exon enhancers across cellular and tissue contexts. 18
Figure S15: PanCancerAtlas variants in exon enhancers and their impact on transcription factor binding. 19
Figure S16: PanCancerAtlas lollipop genomic track. 20
Figure S17: GWAS Catalog Variants in Exonic Enhancers (EEs). 21
Figure S18: Comparative conservation of exonic and intergenic enhancers between human and mice. 22
Figure S19: gnomAD SNP density in exonic enhancers and association with TF binding density. 23
Figure S20: Evolutionary conservation and structural properties of exonic enhancers. 24
Figure S21: Selection of exonic enhancers based on transcription factor summit density. 26
Figure S22: STARR-seq experimental design and validation of exonic enhancer activity. 27
Figure S23: Validation of CRISPRi-competent K-562 cells by inhibition of CD81 expression. 28
Figure S24: Distribution and regulatory impact of gnomAD SNPs in exonic enhancers. 29

**References** 30

### Supplementary tables

##### Table S1: EE tested by reporter luciferase assay Figure 3a.

List of EE genomic regions tested by luciferase assay.

| EE ID | sequences |
|---|---|
| EE1 | chr7:139600417-139600596 |
| EE2 | chr2:33020909-330212 |
| EE3 | chr19:34327990-34328151 |
| EE4 | chr2:85326508-85326804 |
| EE5 | chr20:3744902-3745088 |
| EE6 | chr1:212007250-212007449 |
| EE7 | chr7:105106522-105106772 |
| EE8 | chr10:28120194-28120390 |
| EE9 | chr12:103997451-103997535 |
| EE10 | chr10:27173118-27173259 |
| EE11 | chr14:88472244-88472465 |
| EE12 | chr11:121553937-121554109 |
| EE13 | chr1:29033093-29033245 |
| EE14 | chrX:71394067-71394245 |
| EE15 | chr11:74085618-74085886 |
| EE16 | chr1:113659080-113659265 |
| EE17 | chr19:50414815-50414990 |
| EE18 | chr11:27372283-27372398 |
| EE19 | chr9:129863186-129863299 |
| EE20 | chr3:179375220-179375341 |
| EE21 | chr6:34835294-34835464 |
| EE22 | chr15:41438321-41438431 |
| EE23 | chr6:29631396-29631599 |

##### Table S2: Tested promoters on Figure 3bc.

| | |
|---|---|
| **GARRE1 Promoter (P1 GARRE1)** | chr19:34254350-34254697 |
| **GPI Promoter (P2 GARRE1)** | chr19:34364882-34365266 |
| **Promoter PDCD2L (P3 GARRE1)** | chr19:34404188-34404430 |
| **Promoter USP20 (P1 USP20)** | chr9:129835222-129835554 |
| **Promoter C9orf78 (P2 USP20)** | chr9:129835222-129835554 |
| **Promoter TOR1B (P3 USP20)** | chr9:129802932-129803212 |

##### Table S3: EE mutated on Figure 3de.

Sequences of wt and mutated EEs tested by luciferase

| |
|---|
| EE3_WT |
| GGCTGCTGCAGCGAGGCGGAAGCCCAGCAGACGGGGCGGAGGCAGACACCCCCGCAGCCCATGCAGT<br>GTGAGCTCCCCACCGTCCCTGTGCAGATAGGATCGCACTTCCTGAAGGGCGTCTCCTTTAATGAGTCGG<br>CCGCCGACAATCTGAAACTTAAGACG |
| EE3_gnomaD |
| GGCTGCTGCAGCGAGttaGAAGCCCAGCAGACGGGGaaGAGGCAGACAgCCtCaCAGCCCATGCAGTGTG<br>AGCTCCCCACtaTCCCTGTGCAGATAGGATtcCACTTtCTGAAGGGtGTCTtCTTgAATGAGTtGGCtaCCGACA<br>gTCTGAAACTTAAGAgG |
| EE3_synonymous |
| GGCTGCTGCAGCGAGGCaGAAGCCCAGCAGACGGGGaGGAGGCAGACACCCCCaCAGCCCATGCAGTG<br>TGAGCTCCCCACtGTCCCTGTGCAGATAGGATCcCACTTtCTGAAGGGtGTCTCCTTcAATGAGTCGGCtGC<br>CGACAATCTGAAACTTAAGACG |
| EE3_missense |
| GGCTGCTGCAGCGAGttGGAAGCCCAGCAGACGGGGCaGAGGCAGACAgCCtCGCAGCCCATGCAGTGTG<br>AGCTCCCCACCaTCCCTGTGCAGATAGGATtGCACTTCCTGAAGGGCGTCTtCTTgAATGAGTtGGCCaCCG<br>ACAgTCTGAAACTTAAGAgG |
| EE19_WT |
| CCCGCCGCTGACTCAGTTCTTCTTGGAGTGTGGCGGCCTGGTGCGCACAGATAAGAAGCCAGCCCTGTG<br>CAAGAGCTACCAGAAGCTGGTCTCTGAGGTCTGGCATAAGAAACG |
| EE19_ in silico |
| CCCGCCGCTcACcCAGTTCTTtTTGGAGTGTGGCGGCCTGGTGCGCACgGAcAAGAAGCCAGCCCTGTGC<br>AAGAGCTACCAaAAGCTGGTCTCTGAGGTCTGGCATAAGAAACG |
| EE19_gnomaD |
| CCCGCCGCTGACTtAaTTCTTCTTcGAGTGTGGCGGCCTGGTGCGCACAGggAAGAAGCCAGCCCTGTGC<br>AAGAGCTACCAGAAGCTGGTCTCTGAGGTCTGGCATAAGAAAtG |
| EE20_WT |
| GAGTGTATCTCGCAGTCAGCAGTGAAAACAAAGTTCGAACAGCACACTATCAGAGCTAAACAGATACTAG<br>CTACTGTGAAAAACATAATGGATTCAGTAAACCTGGCAGCTGAAGATAAAAG |
| EE20_in silico |
| GAGTGTATCTCaCAGTCAGCtGTGAAAACAAAGTTgGAACAGCACACTATCAGAGCTAAACAaATACTAGCT<br>ACTGTGAAAAACATAATtGATTCtGTAAACtTGGCAGCTGAAGATAAgAG |

##### Table S4: gRNAs used for CRISPRi Figure 6b-e.

| Name | Sequence |
|---|---|
| GARRE1_gRNA_CRISPRi | CCGACAATCTGAAACTTAAG |
| USP20_gRNA_CRISPRi | CCGCTGACTCAGTTCTTCTT |
| STK11IP_gRNA_CRISPRi | TCCGGAAACCCTCTGCCGGC |
| COG1_gRNA_CRISPRi | TTGACAGATACGCAGATGCG |
| gRNA no target | GGGAACGACTATGACCGCCA |
| CD81_gRNA_CRISPRi | GCCTGGCAGGATGCGCGGTG |

**Table S5: Primers to assess gene expression Figure 6b-e.**

| Gene | Foward | Reverse |
|---|---|---|
| qGARRE1 | CAGCAGTGGAGAGCAAGACAC | CTTTGGGTGGCCACGTTTTG |
| qGARRE1_up ex6 | AAAAATCGACAGTGCTTTGC | CTTTGGATTCAGGTGGAAGC |
| qUSP20 | TCAAAGGAAGCGGCCATGTC | CAGGCTGTTCAGGTAGGTCC |
| qUSP20_up ex9 | CTGAAAGCTGTTCCTATTGC | CAGCGTTCATGTAGCAGGAG |
| qSTK11IP | CGGCTGCTCTTCTACGATG | AACAGCTCCTCCCATGGTTC |
| qSTK11IP_up ex14 | AGCCCCTGCTTCATAAGGTT | GGAGAGGGGTTCAGAGTTCG |
| qCOG1 | ACATCGAAACAAAAGCTCAGGT | GTCTTCTTCACTGACAAGCTGT |
| qSUCO | ACCCCTCAAGTTTTCTCCAG | TGGGGTGCAATGGTTCTTC |
| qPKD1L2 | ATCAGCTGTCGGAGGAAGGG | CAGAGACCCAGGCTGCTC |
| qGPI | TCGTTCAGGGCATCATCTGG | TCACTTGAGCACTGCCATC |
| qC9orf78 | GGCCCTTGAGAGTAGGTGAC | TGCCTTCTCGTTAGCAGGAC |
| qOBSL1 | GTTCGACCTGAGGACCAAGG | CCTAGTTGCCCTCTACCAGC |
| qVCF1 | CAGACTAAGAGGAAACAAAGCGT | CTGTCCGGGCTATTGATGCT |
| qPIGC | CCATTCTACCTCATTCGCTTGC | ACTGAGGAACCTGGACAAGTC |
| qGCSH | CTCTTGCAGAAAATCCAGGACT | AGGGTTACTCAGTGTCATCTTG |
| qPDCD2L | GCTCAAGAGTGCTAATTTAGGTC | GGGGCCAGCAACTCTTCTC |
| qTOR1B | CAGCTGAAGGACCTGGAACC | AGGCAGGAAGGGGATAAAGT |
| qCHPF | CCTGTGCACATGTACCAGCT | TCCCACTGTAACTCCTGGATCT |
| qC17orf80 | CGTCCTGTGTTGTAGCTGGA | TCTCCACTGTACGCCCTTTG |
| qCD81 | TTCTCCGGGAAGCTGTACCT | ACCATGCTCAGGATCATCTCG |

### Supplementary figures

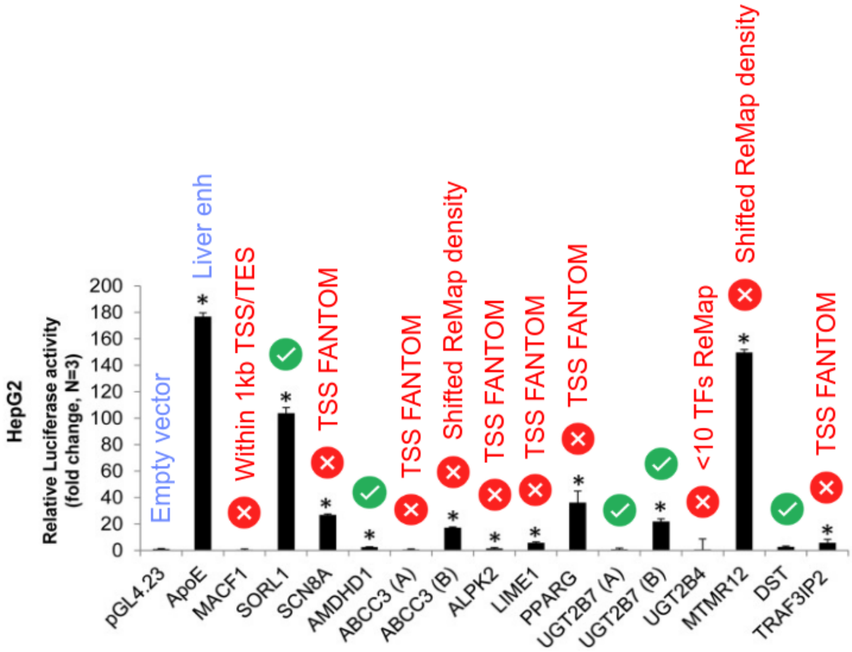


**Figure S1: Re-evaluation of exon enhancer candidates from Birnbaum et al.**

Barplot representing a re-analysis of enhancer activity for 15 exon candidates originally tested in HepG2 cells, as reported by Birnbaum et al.[1]. Exons were lifted to hg38 assembly and filtered based on stringent exon enhancer (EE) criteria. Excluded exons are labelled in red along with the exclusion parameters, including proximity to transcription start site (TSS) or transcription end site (TES), overlap with FANTOM TSS, shifted ReMap ChIP-seq peaks density, or insufficient TF ChIP-seq peaks (<10 TFs in ReMap). The y-axis represents relative luciferase activity in HepG2 cells (fold change, N≥3). Black bars indicate enhancer activity levels, with significant changes marked by asterisks. The control empty vector and a known liver enhancer are included as references. This figure is adapted from Birnbaum et al. [1].

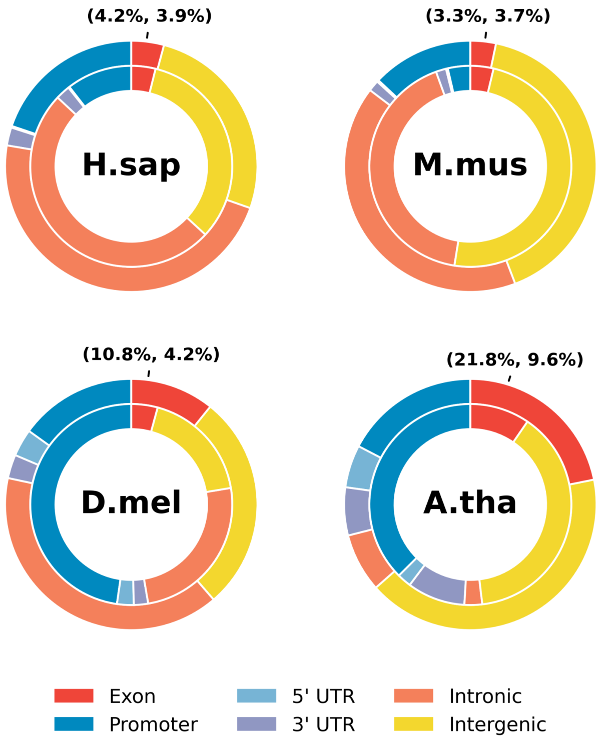


**Figure S2: Genomic distribution of ChIP-seq and DNase-seq data across four species.**

The outer layer represents the distribution of non-redundant transcription factor summits from ReMap2022 ChIP-seq data across *Homo sapiens* (*H.sap*), *Mus musculus* (*M.mus*), *Drosophila melanogaster* (*D.mel*), and *Arabidopsis thaliana* (*A.tha*), annotated using ChIPseeker v.3.20[2]. The inner layer shows the distribution of DNase-seq data obtained from Meuleman et al.[3] for *H.sap*, *ENCODE* for *M.mus*, *ChIP-Atlas* for *D.mel*, and *PlantRegMap* for *A.tha*. The percentages above each species correspond to the proportion of TF ChIP-seq (outer) and DNase-seq sites (inner) located in exonic regions (e.g. For M.mus, 3.3% of ReMap ChIP-seq and 3.7% of DNAseI in coding exons).

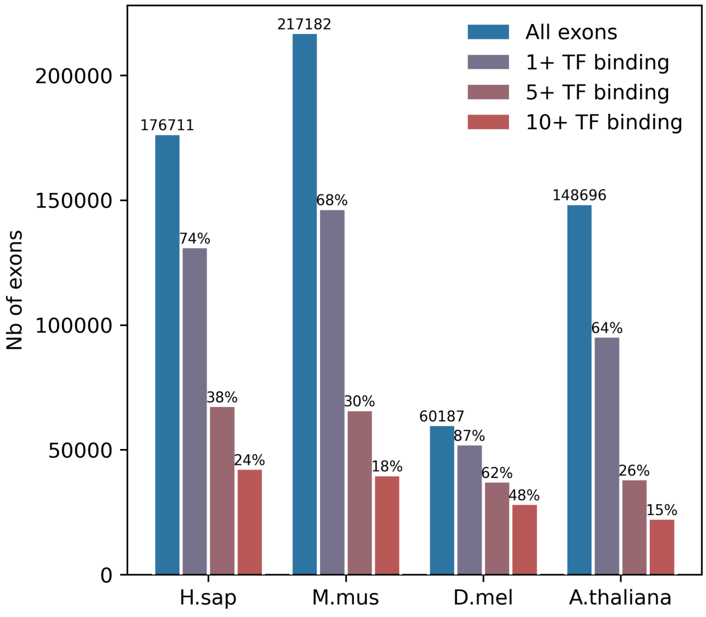


**Figure S3: Transcription factor occupancy in coding exons across species.**

Bar plot showing the number of merged protein-coding exons overlapping non-redundant transcription factor (TF) ChIP-seq summits from ReMap2022 across four species: Homo sapiens (H.sap), Mus musculus (M.mus), Drosophila melanogaster (D.mel), and Arabidopsis thaliana (A.tha). Bars indicate the total number of exons (blue) and subsets with different degrees of TF binding: ≥1 TF (grey), ≥5 TFs (brown), and ≥10 TFs (red). Percentages above each bar represent the proportion of exons in each binding category relative to the total number of exons for that species. Total exon counts are displayed above the blue bars for each species.

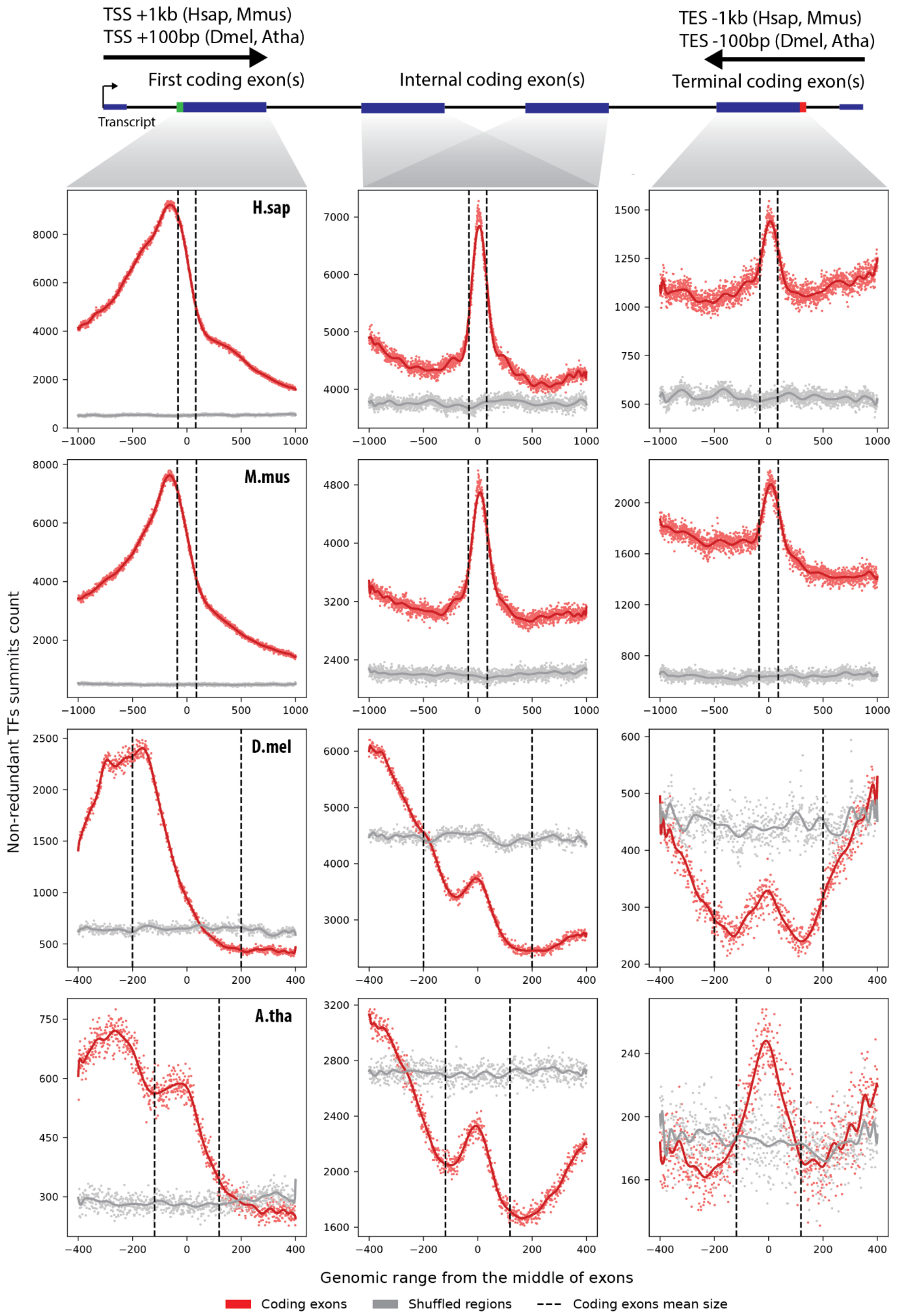


**Figure S4: Genomic distribution of transcription factor binding across exon classes.**

Meta-profile analysis showing genomic distribution of non-redundant transcription factor (TF) ChIP-seq summits from ReMap2022 over merged protein-coding exons across four species: *Homo sapiens* (*H.sap*), *Mus musculus* (*M.mus*), *Drosophila melanogaster* (*D.mel*), and *Arabidopsis thaliana* (*A.tha*). The top schematic categorizes exons into three regions: transcription start site (TSS)-proximal exons (±1000 bp for *H.sap* and *M.mus*; ±100 bp for *D.mel* and *A.tha*), internal coding exons, and transcription end site (TES)-proximal exons (±1000 bp for *H.sap* and *M.mus*; ±100 bp for *D.mel* and *A.tha*). The line plots represent the distribution of TF ChIP-seq summits across exonic regions in each category and each species. TF occupancy in coding exons is shown in red, while shuffled control regions are represented in grey. Dashed vertical lines indicate mean exon boundaries. This analysis reveals TF-binding enrichment patterns along exons.

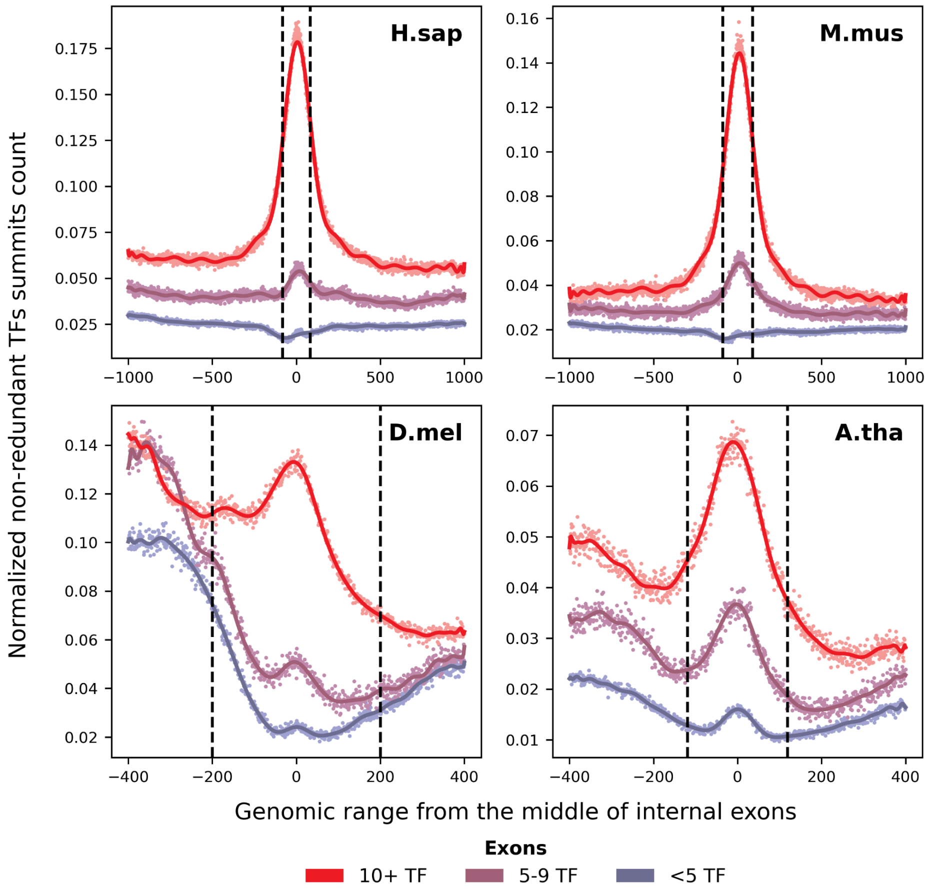


**Figure S5: Transcription factor binding density in internal exons.**

Meta-profile analysis showing the normalized distribution of non-redundant transcription factor (TF) ChIP-seq summits from ReMap2022 in internal exons of Homo sapiens (H.sap), Mus musculus (M.mus), Drosophila melanogaster (D.mel), and Arabidopsis thaliana (A.tha). Data are stratified by TF binding density: exons bound by <5 TFs (blue), 5-9 TFs (purple), and ≥10 TFs (red). The x-axis represents the genomic distance (bp) from the center of internal exons, with dashed vertical lines indicating average exon boundaries for each species. The y-axis represents the normalized frequency of TF ChIP-seq summits. Lines show the smoothed average profile, while dots represent individual data points. The data reveal a clustering of TF binding around exon centers, with a stronger enrichment in exons containing a higher number of TF peaks, highlighting potential.

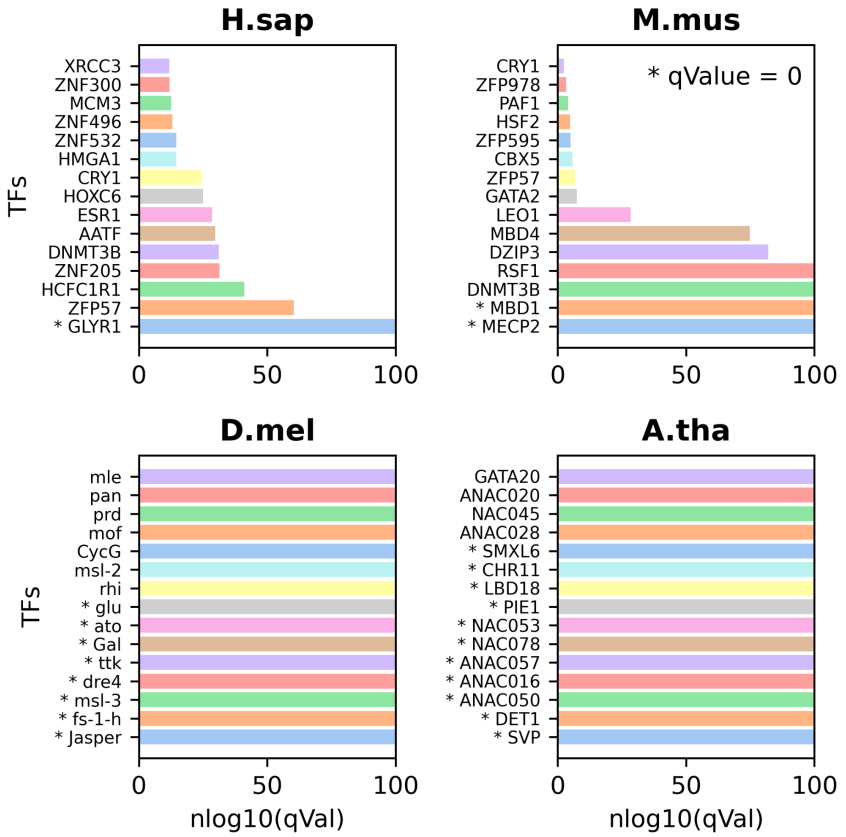


**Figure S6: Enrichment of transcription factors in EEs**

Enrichment analysis of transcription factors (TFs) binding to exonic enhancers (EEs) was performed using non-redundant ReMap2022 ChIP-seq datasets across four species: *Homo sapiens* (*H.sap*), *Mus musculus* (*M.mus*), *Drosophila melanogaster* (*D.mel*), and *Arabidopsis thaliana* (*A.tha*). Enrichment was assessed using LOLA[4] (v.1.12.0) by comparing EE-associated TFs to promoter-associated TFs from the Eukaryotic Promoter Database (EPD, https://epd.expasy.org/epd/).

The top 15 enriched TFs in each species are ranked by -log10(q-value), with significantly enriched TFs marked with an asterisk. *H.sap* and *M.mus* show strong enrichment for ZNF-family proteins, chromatin remodelers, and DNA methylation regulators, while *D.mel* and *A.tha* exhibit enrichment for species-specific transcription factors involved in developmental regulation. These results highlight the distinct TF repertoires associated with EEs across species.

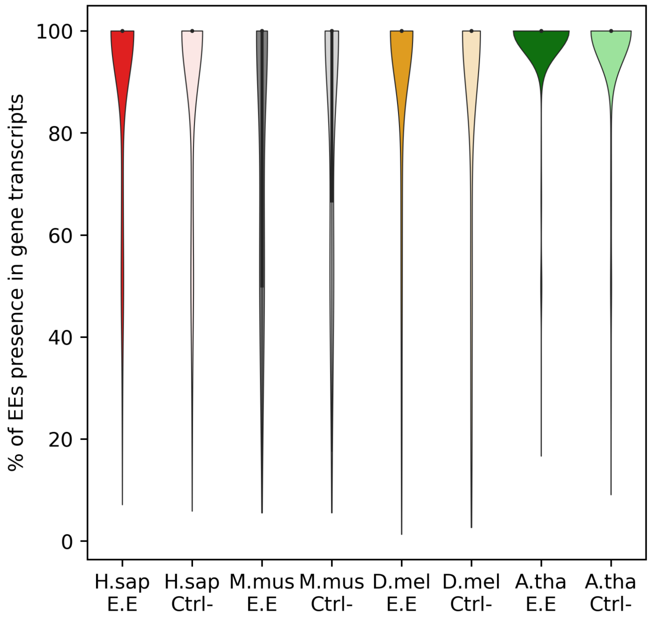


**Figure S7: Prevalence of exon enhancers across gene transcript isoforms.**

EEs were mapped to all protein-coding exons, and their representation in transcripts was assessed by calculating the proportion of transcripts containing the EE relative to the total number of transcripts per gene. Violin plots show the distribution of this proportion across four species: Homo sapiens (H.sap), Mus musculus (M.mus), Drosophila melanogaster (D.mel), and Arabidopsis thaliana (A.tha). Control exons (Ctrl−) are included for comparison. In all species, EEs are present in a substantial fraction of gene transcripts, indicating their widespread inclusion in alternative isoforms. These results suggest a potential regulatory role for EEs in transcript diversity and gene expression modulation.

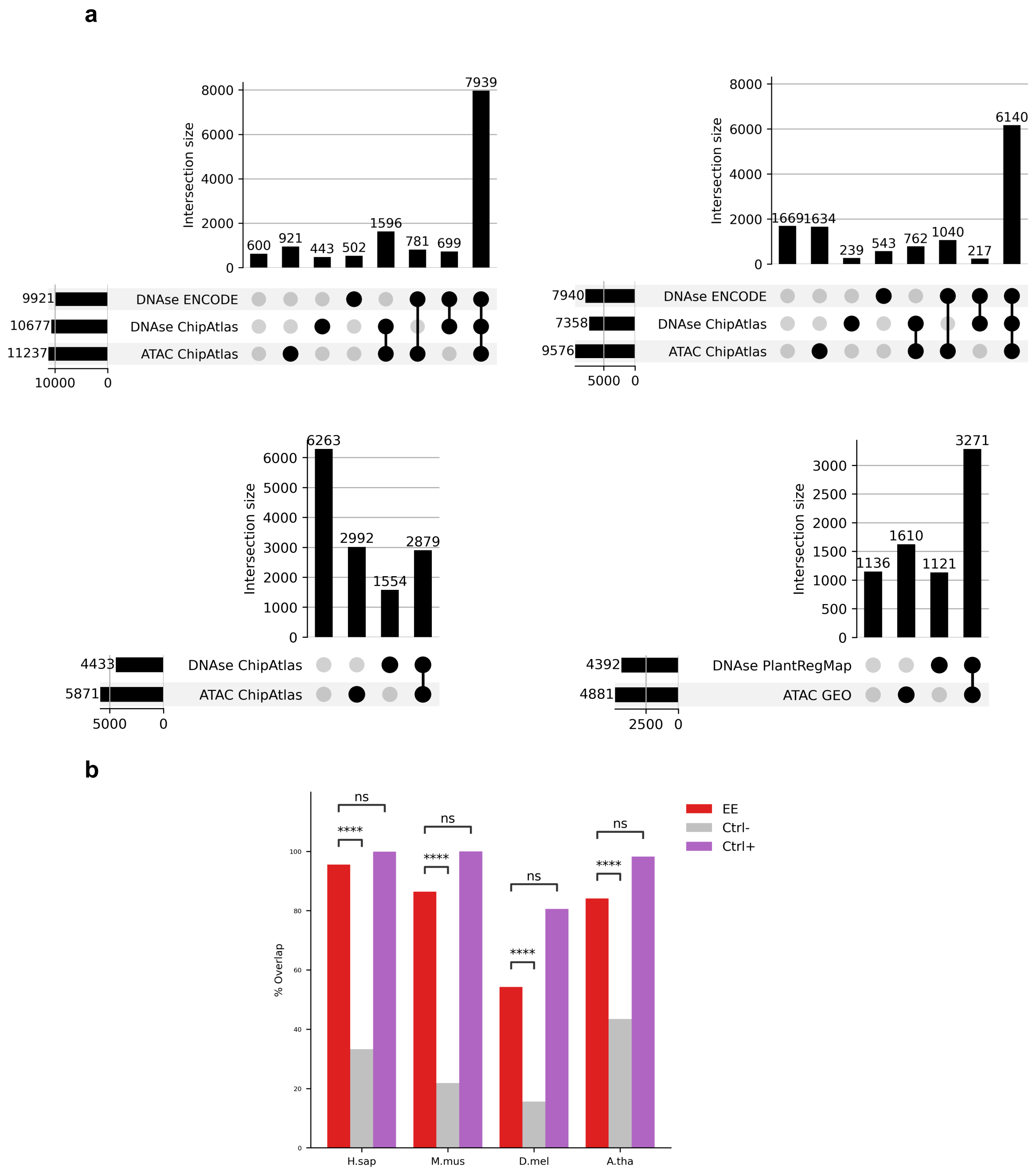


**Figure S8: Chromatin accessibility at exon enhancers across species.**

(**a**) Overlap of EEs with DNase-seq and ATAC-seq peaks from ENCODE, ChipAtlas, PlantRegMap, and GEO datasets. UpSet plots depict the intersection sizes of EEs with chromatin accessibility datasets, highlighting the fraction of EEs that overlap multiple sources. (**b**) Quantification of DNase-seq and ATAC-seq peak overlap in EEs, control exons with no TF binding (Ctrl−), and highly bound control exons (Ctrl+). Fisher's exact one-side test (****P < 0.00005) was used to assess statistical differences in chromatin accessibility between EEs and control exons.

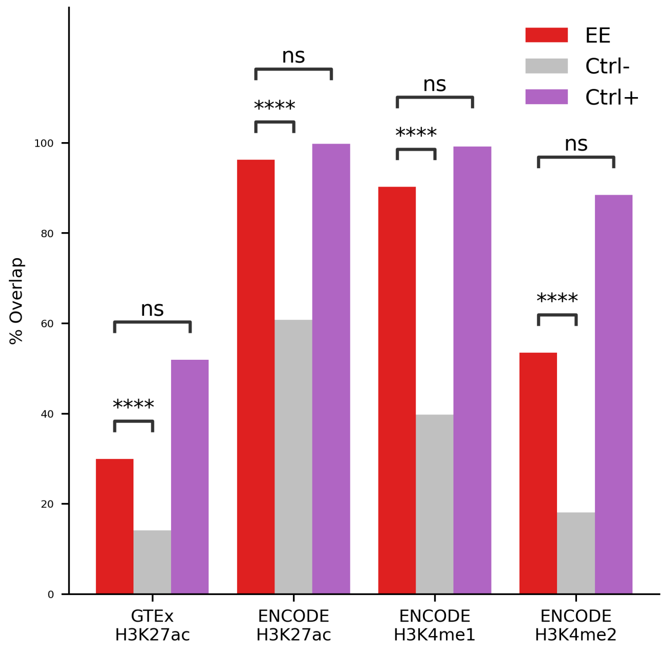


**Figure S9: Enrichment of histone modifications at exon enhancers.**

Histone modification enrichment at exon enhancers (EEs) in H. sapiens. Overlap of EEs with H3K27ac marks from GTEx (5 tissues) and ENCODE (853 datasets), as well as with H3K4me1 (388 datasets) and H3K4me2 (77 datasets) from ENCODE. The proportion of EEs overlapping histone modifications was compared to control exons with no TF binding (Ctrl−) and highly bound control exons (Ctrl+). Statistical significance of differences in overlap between EEs and controls was assessed using Fisher's exact one-side test (****P < 0.00005).

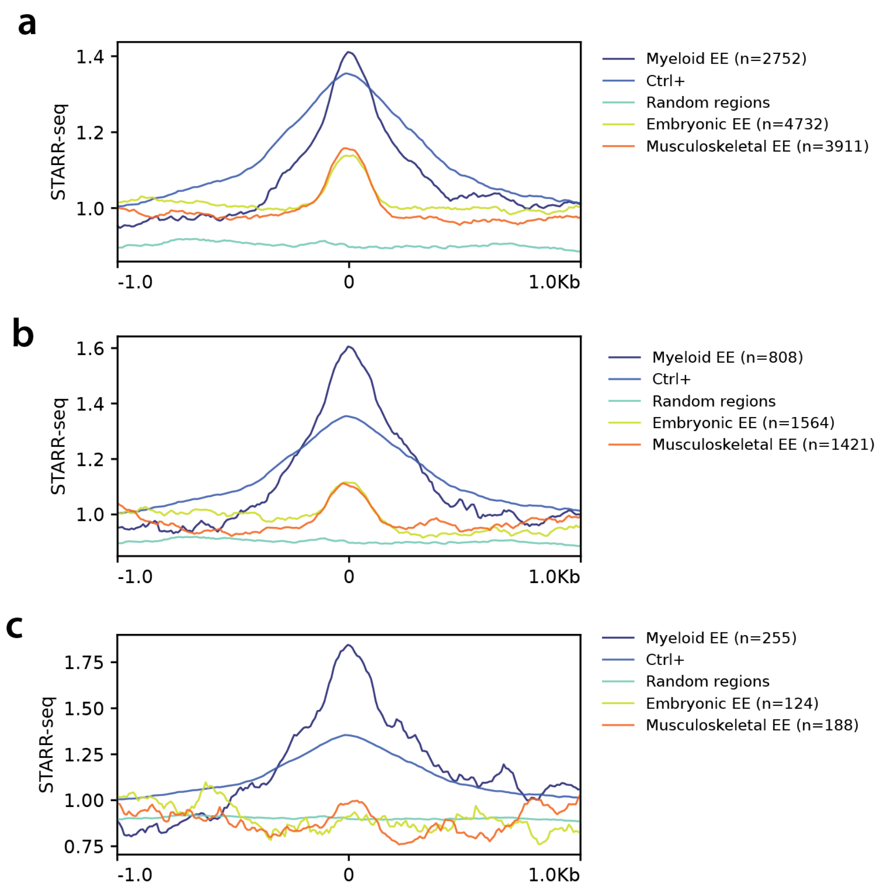


**Figure S10: STARR-seq activity in exonic enhancers stratified by biotype**

STARR-seq signal intensity for exonic enhancers (EEs) in K-562 cells, categorized based on biotype specificity. **(a)** EEs where myeloid TF ChIP-seq peaks are among the top three most frequent biotypes. **(b)** EEs where myeloid TF ChIP-seq peaks represent the predominant biotype. **(c)** EEs where myeloid TF ChIP-seq peaks are the predominant biotype, with at least 50% of EE-associated transcription factors (TFs) linked to that biotype. The data show that STARR-seq activity increases with myeloid-specific TF occupancy, with the strongest signal observed in EEs enriched for myeloid TF ChIP-seq peaks. In contrast, STARR-seq activity progressively decreases in EEs associated with embryonic and musculoskeletal biotypes, indicating that enhancer activity is shaped by cell-type-specific transcription factor binding.

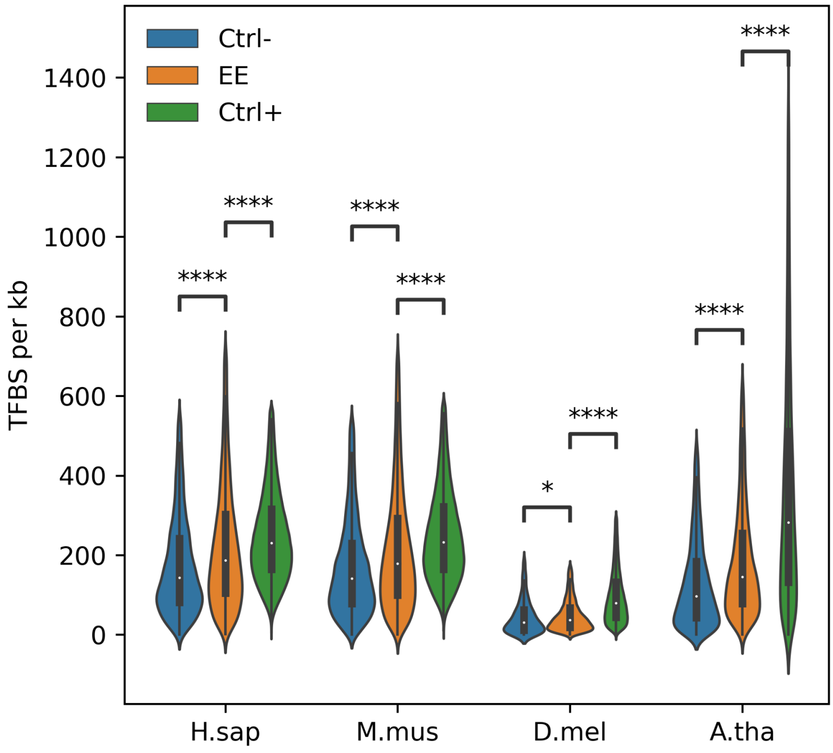


**Figure S11: Transcription factor binding site density at exon enhancers.**

Transcription factor binding site (TFBS) density per kilobase at exon enhancers (EEs) across four species. TFBSs were predicted using JASPAR2024 with a threshold score ≥ 400 and overlapped with EEs, as well as control exons with no TF binding (Ctrl−) and highly bound control exons (Ctrl+). Statistical significance was assessed using a Student's t-test ($P < 0.05$). Enhancers (Ctrl+) and exon enhancers contain significantly more in silico predicted TFBSs than Ctrl− sequences, consistently across all species, suggesting a conserved enrichment of regulatory sequence features within these elements.

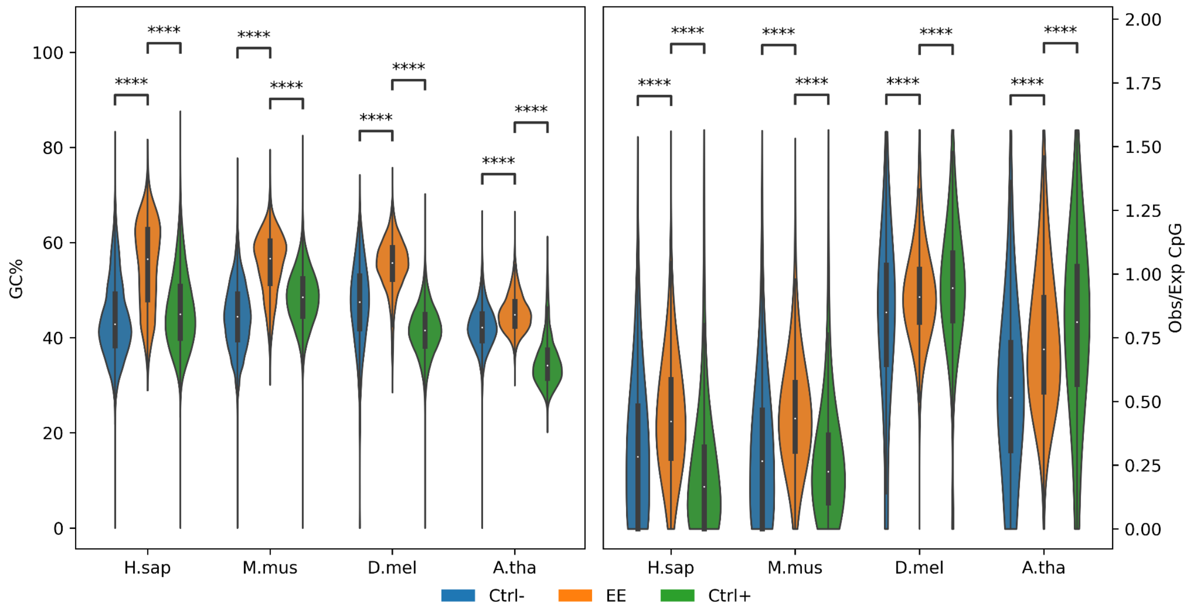


**Figure S12: GC content and CpG enrichment at exon enhancers.**

Comparative analysis of GC content and CpG dinucleotide frequency at exon enhancers (EEs) across four species. The left panel shows the GC content of EEs compared to control exons with no TF binding (Ctrl−) and highly bound control exons (Ctrl+). The right panel presents the observed-to-expected CpG ratio ((CG)/((CxG)/L)) for EEs and controls. Statistical significance was assessed using a Student's t-test ($P < 0.05$), highlighting a distinct GC and CpG profile in EEs.

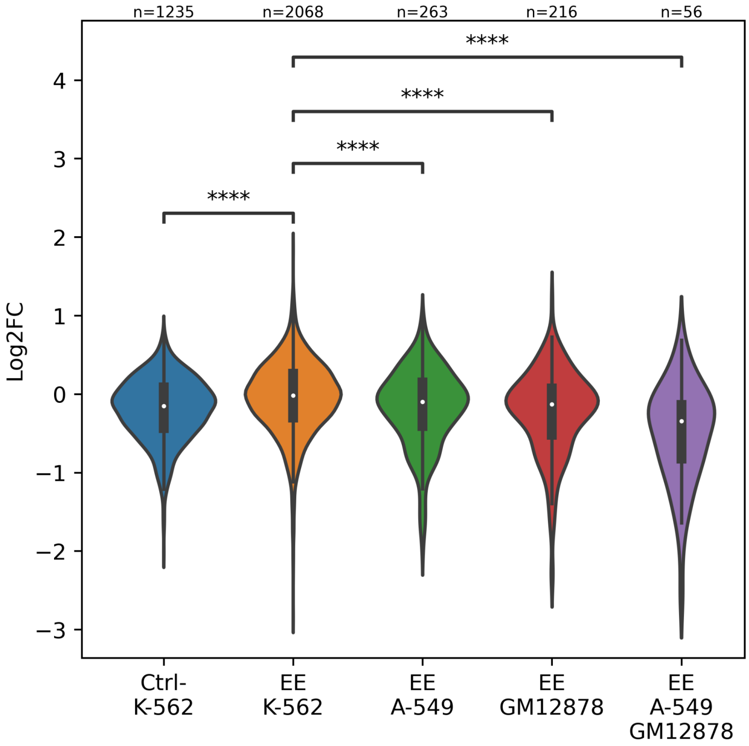


**Figure S13: Cell line specificity and selection refinement of exon enhancer activity in STARR-seq.**

Exon enhancers (EEs) exhibit significant differential activity across cell lines and selection criteria in STARR-seq assays. EEs selected based on associated A-549 or GM12878 cell line signatures show significantly lower activity in K-562 cells Statistical significance was assessed using a *t*-test (*****P* < 0.00001).), suggesting context-dependent regulatory function.

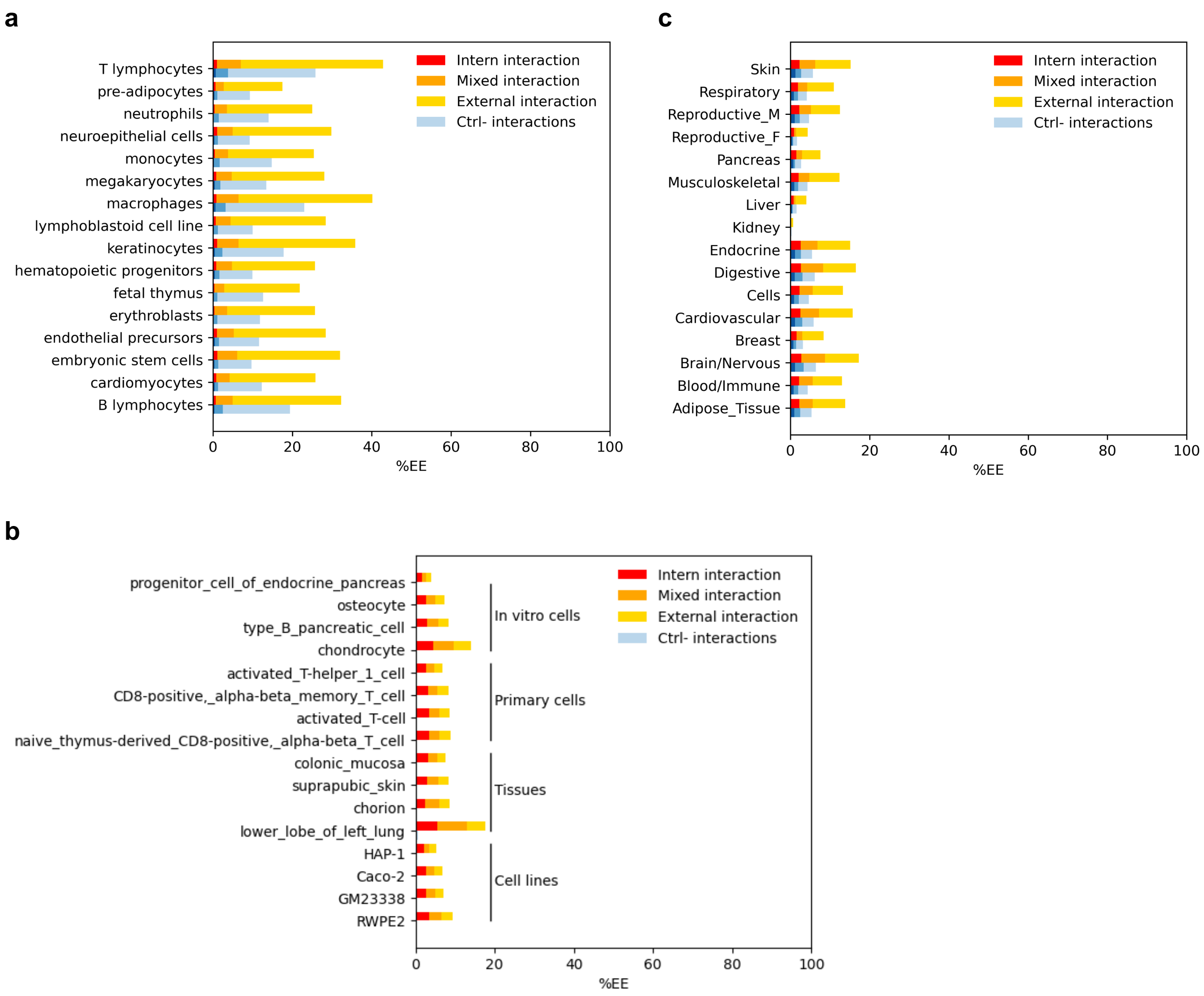


**Figure S14: Interaction landscape of exon enhancers across cellular and tissue contexts.**

Exon enhancers (EEs) exhibit distinct chromatin interactions with target genes, categorized as internal (within the host gene), external (distal genes), or mixed (host + distal interactions). **(a)** Normalized pCap-HiC interactions from Laverré et al.[5] reveal the prevalence of these interaction types across diverse human cell lines, highlighting variations in EE connectivity depending on the cellular context. **(b)** ENCODE-E2G interaction data show the four most frequent EE interactions, classified into *in vitro* cells, primary cells, tissues, and cell lines. **(c)** GTEx eQTL associations link EEs to their target genes across 54 human tissues, normalized into 16 biotypes following the framework of De Langen et al. (*Cell Genom.* 2023).

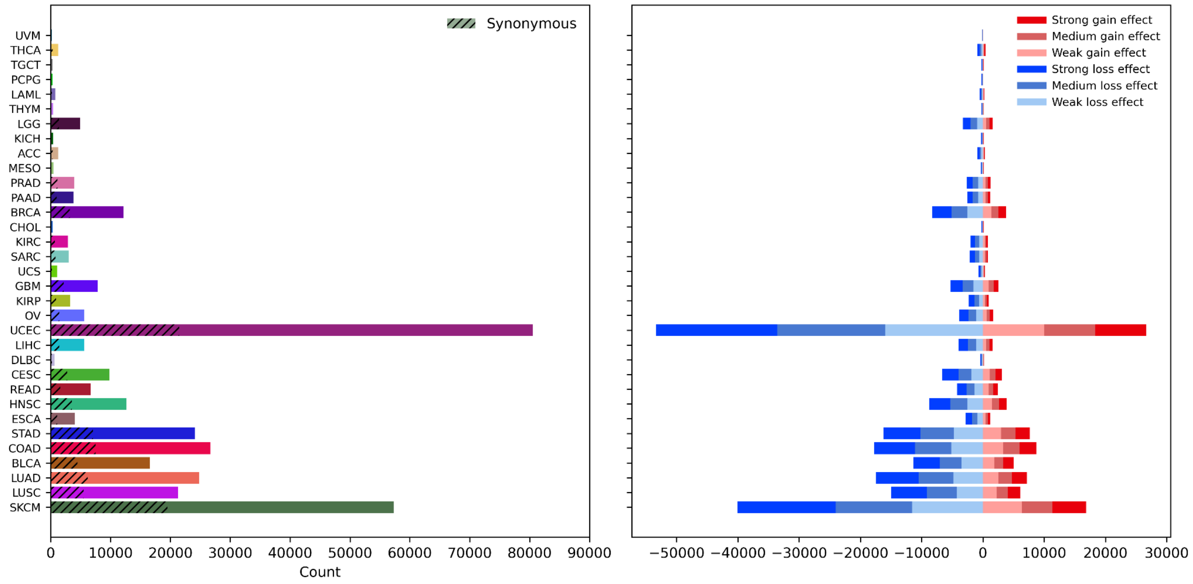


**Figure S15: PanCancerAtlas variants in exon enhancers and their impact on transcription factor binding.**

Distribution and predicted regulatory impact of single nucleotide polymorphisms (SNPs) from the PanCancerAtlas overlapping exon enhancers (EEs). SNPs were mapped to the hg19 reference genome using UCSC liftOver. Left panel: The number of unique SNPs identified in EEs for each cancer type. Hatched bars represent synonymous variants, while coloured bars denote different cancer types. Right panel: Predicted effect of SNPs on transcription factor binding sites (TFBSs) assessed using the FABIAN-variant tool with JASPAR motifs, categorized into strong (dark), medium, and weak (light) effects for TF binding gain (red) and TF binding loss (blue). This analysis suggests that somatic mutations in EEs may alter transcription factor binding, potentially contributing to regulatory changes across different cancer types.

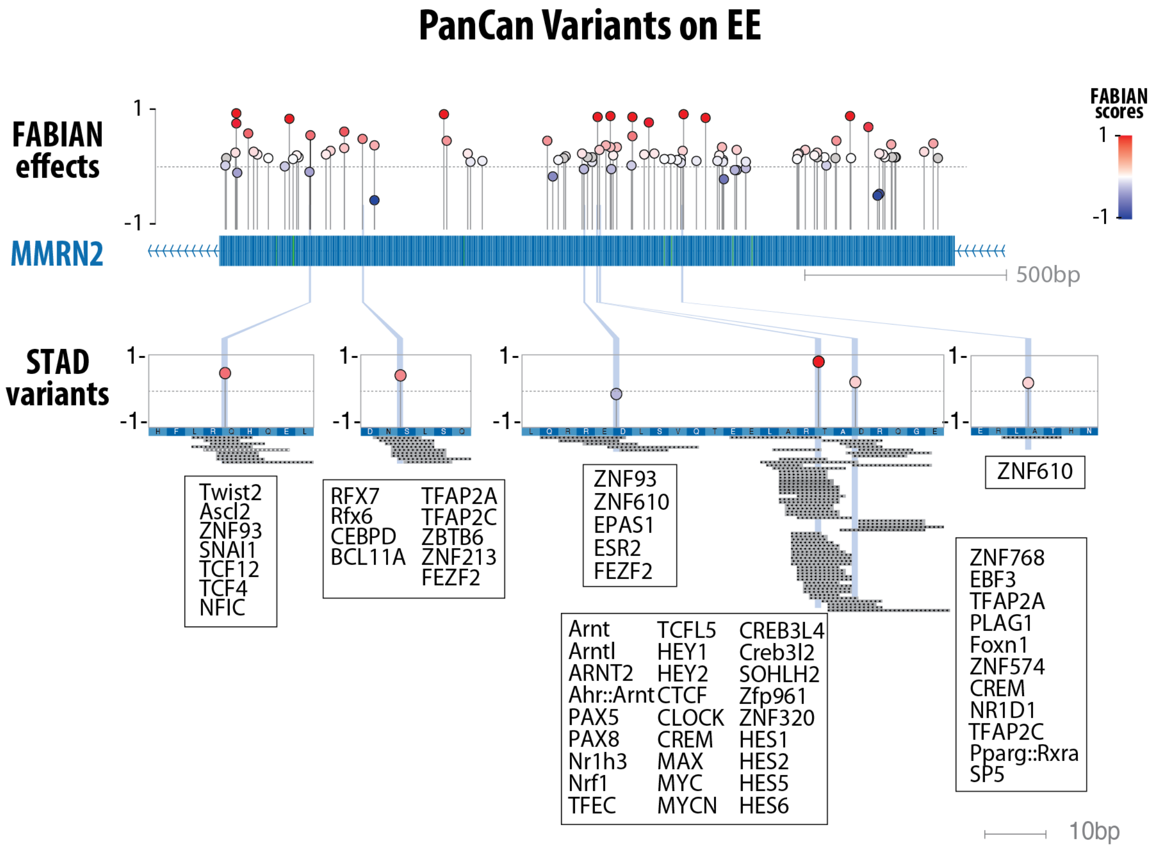


**Figure S16: PanCancerAtlas lollipop genomic track.**

Example genomic track at the *MMRN2* locus, illustrating predicted transcription factor binding changes from FABIAN variant analysis (4). Red lollipops indicate variants that increase TF binding affinity, whereas blue lollipops denote decreased affinity. The *STAD* (gastric cancer) variants align with multiple JASPAR TF motif sites (listed).

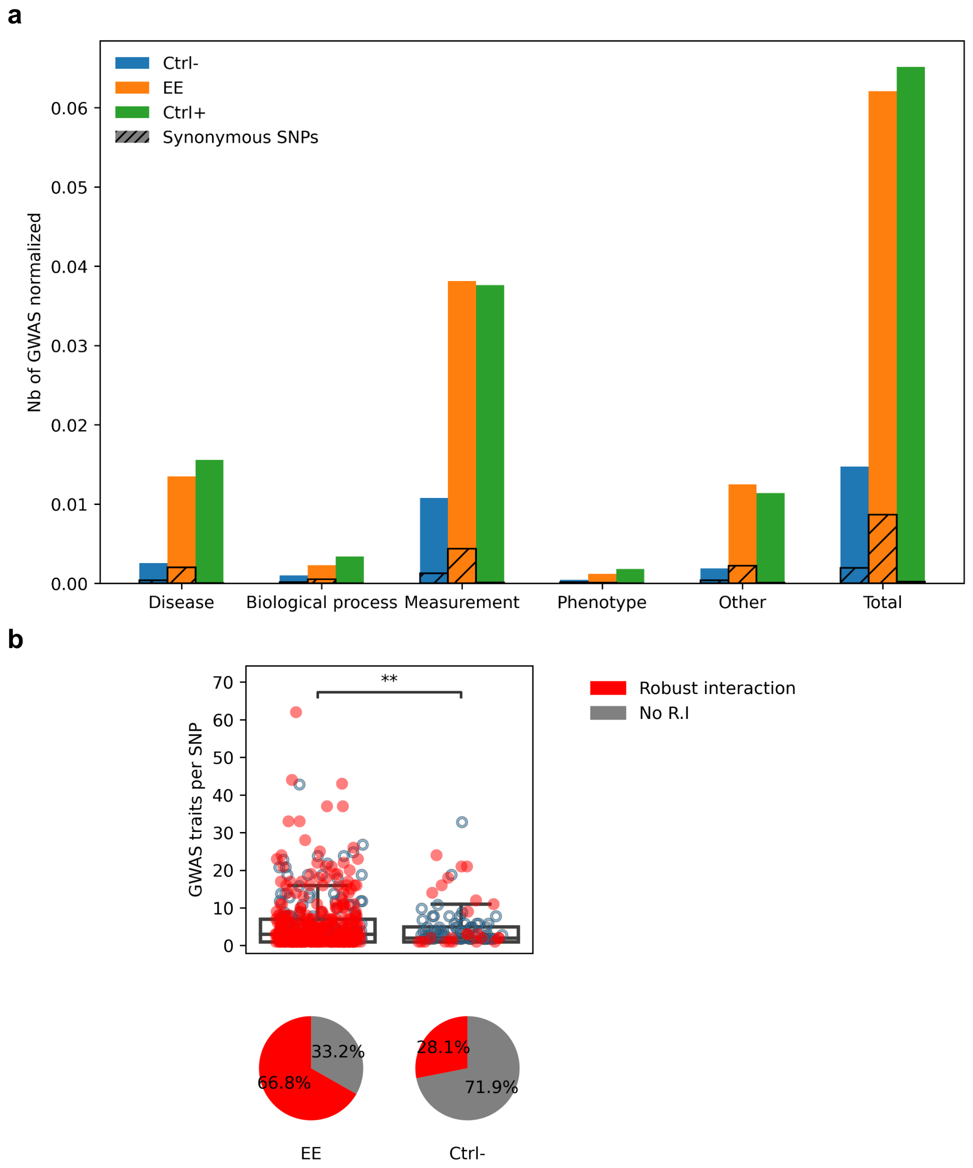


**Figure S17: GWAS Catalog Variants in Exonic Enhancers (EEs).**

**(a)** Overlap of GWAS-SNPs (linkage disequilibrium r2 > 0.8) with EEs. GWAS traits were mapped into five parent traits (i.e Disease, Biological process, Measurement, Phenotype and Other). For each category, the number of GWAS-SNPs overlapped by EEs (total=837) and normalized by the dataset size (EE=13,481; Ctrl-=13,253; Ctrl+=404,325) shows a significantly higher overlap compared to negative controls (total=195) and overlap rates similar to positive control (n=26,347). Moreover, we denote the presence of synonymous GWAS-SNPs overlapping EEs in all categories.

**(b)** Overlap of pleiotropic GWAS SNPs retrieved from Watanabe et al., with EEs. In red are SNPs in exon with a robust interaction coming from the crossover of pCap-HiC, ENCODE-rE2G and GTEx eQTLs data. The difference in the numbers of traits associated per SNP between the EE and control dataset was assessed with an Anderson-Darling test (P=0.0025).

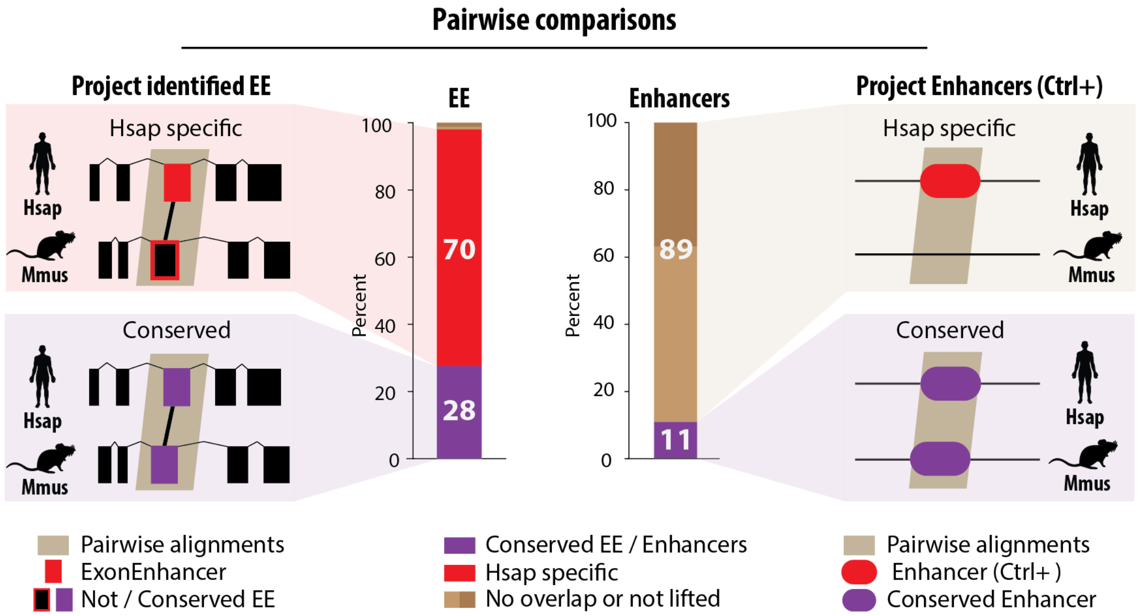


**Figure S18: Comparative conservation of exonic and intergenic enhancers between human and mice.**

Pairwise conservation of exonic enhancers (EEs) and intergenic enhancers (Ctrl+) between *Homo sapiens* (*H.sap*) and *Mus musculus* (*M.mus*) using LastZ-net pairwise alignments. **Left panel (EEs):** The proportion of EEs classified as *H.sap*-specific (70%) versus conserved between both species (28%). **Right panel (Enhancers):** Comparison with ENCODE intergenic enhancers used as positive controls, showing a lower conservation rate (11%) compared to EEs (28%). Annotations; Red: *H.sap*-specific EEs or enhancers. Purple: Conserved EEs or enhancers. Beige: Pairwise alignments between species. These results suggest that a larger fraction of EEs are evolutionarily conserved compared to classical intergenic enhancers, highlighting their potential functional significance.

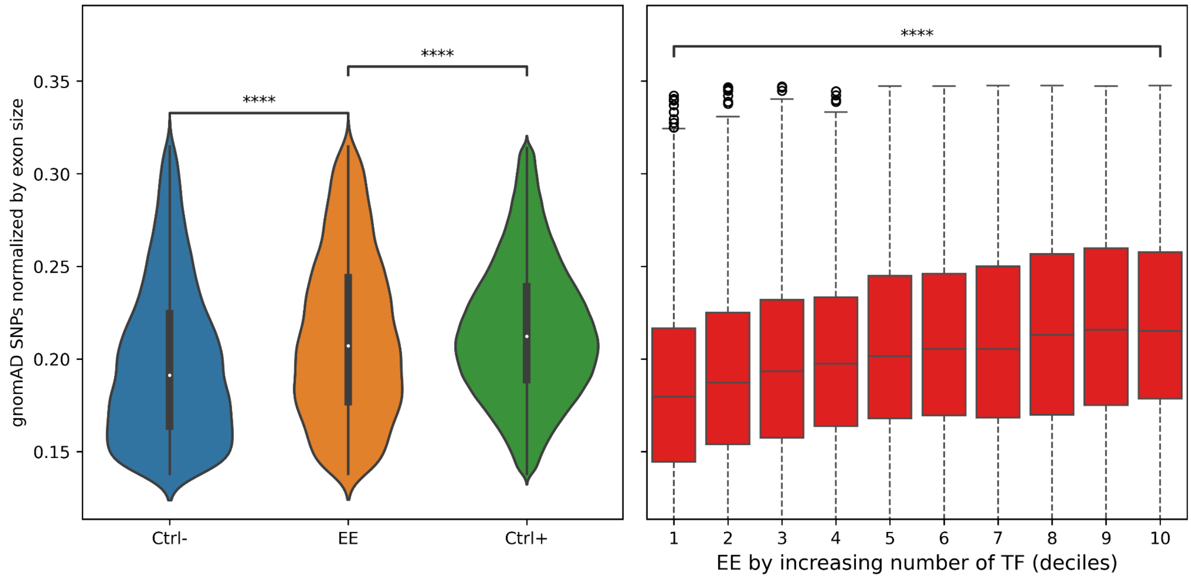


**Figure S19: gnomAD SNP density in exonic enhancers and association with TF binding density.**

Distribution of gnomAD v.3 SNPs within exonic enhancers (EEs), normalized by exon size, compared to control datasets. Left panel: SNP density comparison between EEs (orange), negative control exons (Ctrl−, blue), and intergenic enhancers (Ctrl+, green). Statistical significance was assessed using a Student's t-test (****P < 0.00001). Right panel: SNP density in EEs grouped by transcription factor (TF) binding density, ranked by deciles. Statistical significance was evaluated using a Kruskal-Wallis test (****P < 0.00001). Outliers were removed for clarity. These results suggest that EEs have a significantly higher SNP density compared to negative controls and show an increasing trend with TF binding density, indicating potential regulatory constraint.

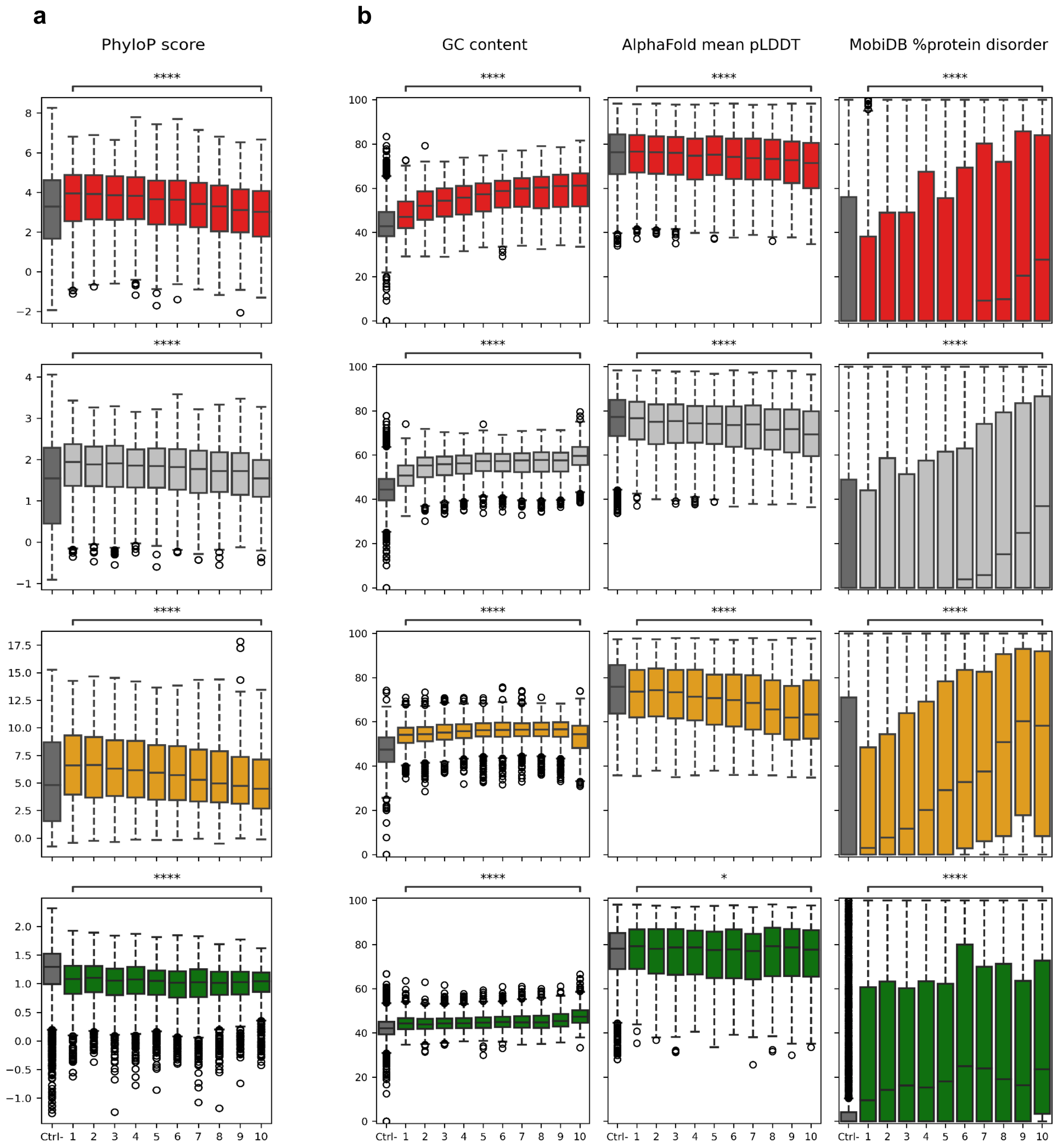


**Figure S20: Evolutionary conservation and structural properties of exonic enhancers.**

Conservation and structural characteristics of exonic enhancers (EEs), grouped by transcription factor (TF) binding density (deciles) across four species: *Homo sapiens* (*H.sap*), *Mus musculus* (*M.mus*), *Drosophila melanogaster* (*D.mel*), and *Arabidopsis thaliana* (*A.tha*).

**(a)** Evolutionary conservation: PhyloP scores represent evolutionary constraints across species, with higher scores indicating stronger conservation. PhyloP scores are derived from multiple genome alignments: 100 species for *H.sap*, 35 species for *M.mus*, 124 species for *D.mel*, and 63 species for *A.tha*.

**(b)** Structural characteristics of EEs:

- GC content: The proportion of guanine-cytosine nucleotides in EE sequences, analyzed across TF-binding deciles.
- AlphaFold mean pLDDT scores: Predicted protein structure confidence scores for EEs, with higher values reflecting more stable structural domains.
- MobiDB % protein disorder: Proportion of intrinsically disordered protein regions in EEs, suggesting increased flexibility in TF-bound exons.

All statistical comparisons were conducted using the Kruskal-Wallis test (****$P < 0.00001$), revealing significant differences in evolutionary conservation, nucleotide composition, and structural properties of EEs across TF-binding densities.

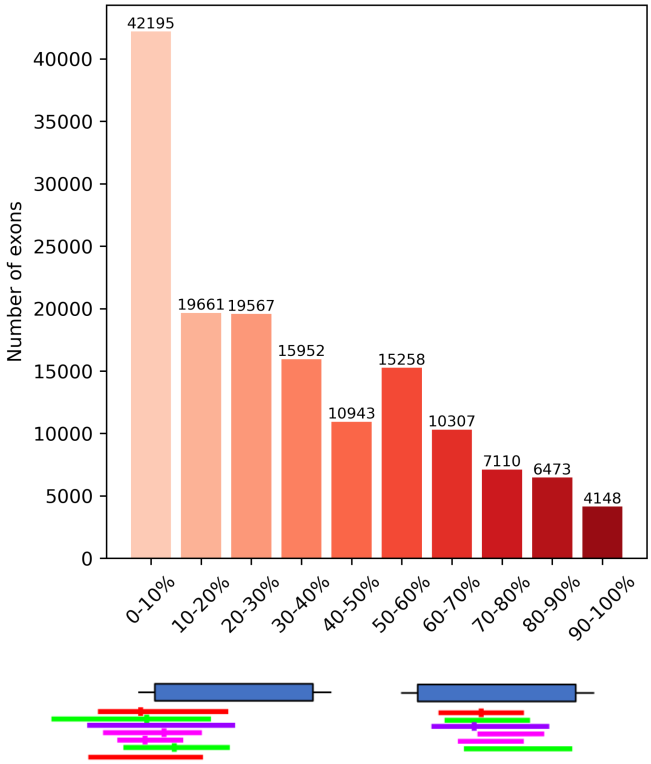


**Figure S21: Selection of exonic enhancers based on transcription factor summit density.**

Distribution of transcription factor (TF) ChIP-seq summits relative to TF number in *Homo sapiens* merged protein-coding exons. Top panel: Histogram displaying the number of exons (y-axis) categorized by TF ChIP-seq summit-to-TF number ratios (x-axis) in percentage bins. Most exons fall within the 0–10% range, with a progressively lower number of exons observed in higher TF ChIP-seq summit ratio bins. Bottom panel: Schema visualization of TF ChIP-seq summit distributions across selected exons, illustrating variance and thresholds for defining exonic enhancers (EEs). To ensure stringent selection criteria, only exons with a TF ChIP-seq summit-to-TF number ratio greater than 50 were retained for EE classification.

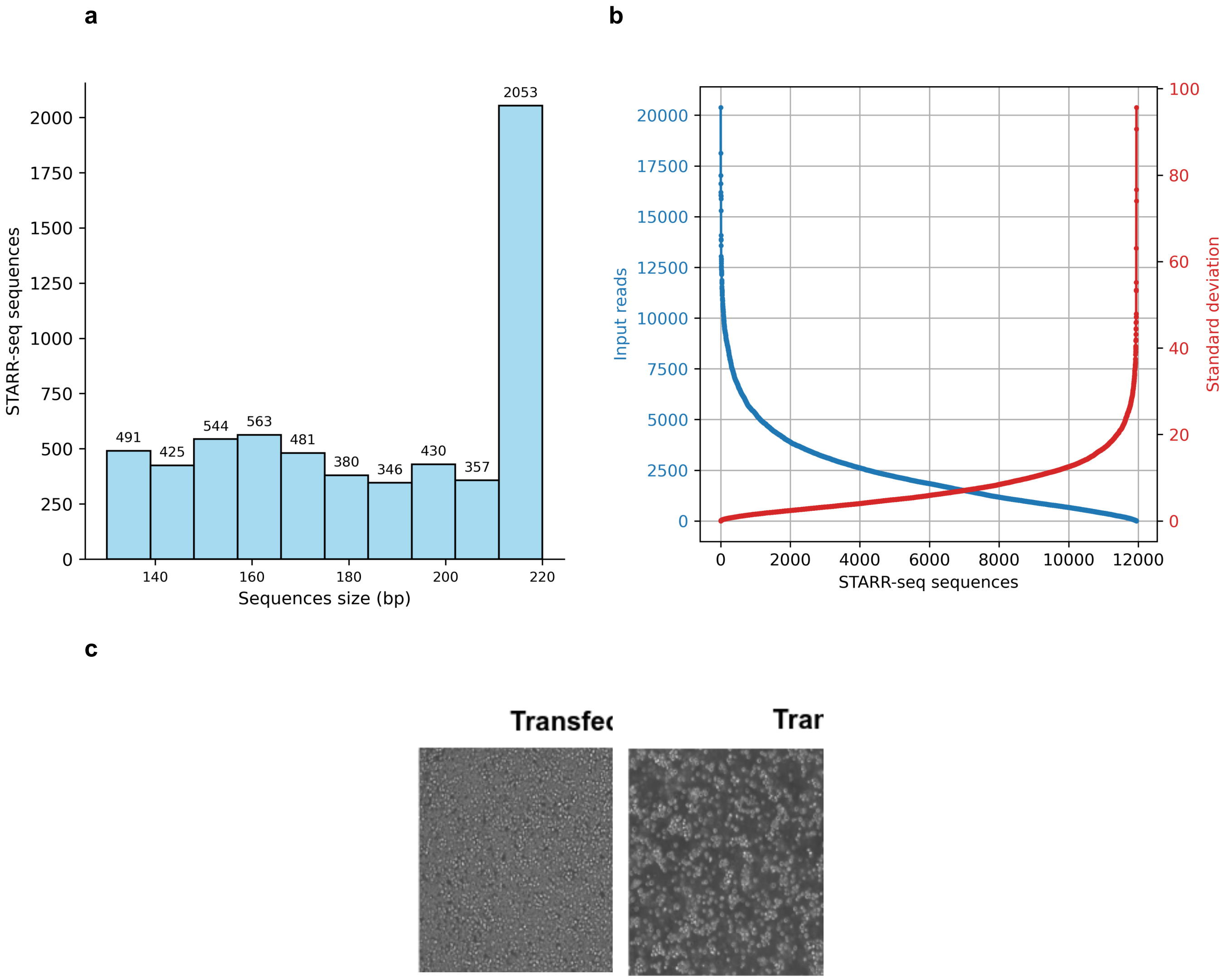


**Figure S22: STARR-seq experimental design and validation of exonic enhancer activity.**

Overview of the STARR-seq experimental setup, including sequence characteristics, input read distributions, and transfection results. **(a) Sequence size distribution:** Histogram showing the distribution of tested sequence lengths (in base pairs, bp). Most sequences are between 140–220 bp, with a peak at 220 bp. **(b) Input read distribution and standard deviation:** The number of input reads (blue curve) and the standard deviation (red curve) across STARR-seq sequences. **(c) Transfection efficiency:** Fluorescence microscopy images comparing cells transfected with empty STARR-seq vector (left) versus the exonic enhancer STARR-seq library (right). The transfected library shows a significantly higher number of fluorescent cells, indicating successful expression from enhancer-active sequences. These data validate the selection and functional testing of candidate exonic enhancers using STARR-seq.

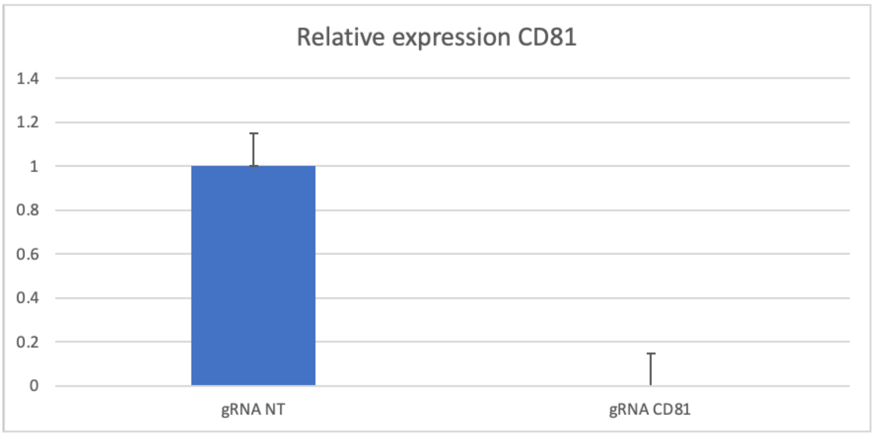


**Figure S23: Validation of CRISPRi-competent K-562 cells by inhibition of CD81 expression.**

Validation of CRISPRi-mediated in K-562 cells stably expressing dCas9–KRAB–MeCP2 was performed with specific guide RNA targeting the CD81 promoter (gRNA CD81) and selected with puromycin. CD81 expression was measured by RT-qPCR and is presented relative to the non-targeting control. The marked reduction in CD81 mRNA confirms efficient CRISPRi knockdown in the dCas9–KRAB–MeCP2 cell line.

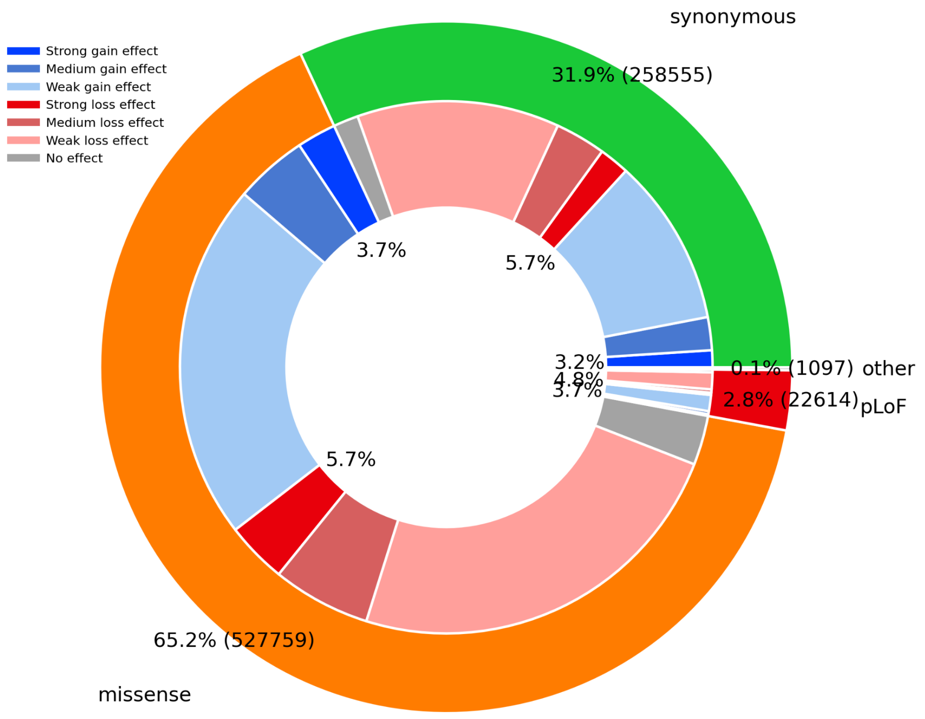


**Figure S24: Distribution and regulatory impact of gnomAD SNPs in exonic enhancers.**

This figure illustrates the distribution of gnomAD v.3 SNPs within exonic enhancers (EEs) and their predicted effects on transcription factor (TF) binding, as assessed by the FABIAN-variant tool. The analysis considers SNPs overlapping a ReMap TF peak and its corresponding transcription factor binding site (TFBS) from JASPAR.

Outer ring: Proportion of SNP types found in EEs, including synonymous (green, 31.9%), missense (orange, 65.2%), pLoF (predicted loss-of-function, red, 2.8%), and other variants (grey, 0.1%).

Inner ring: TF-binding disruption effects of SNPs, with blue shades indicating gain of TF binding and red shades indicating loss of TF binding. Each effect is further classified into three categories based on disruption scores: Strong (>0.66, dark shades), Medium (0.33–0.66, mid shades), and Weak (>0 but <0.33, light shades). Gray represents SNPs with no TF/TFBS overlap.

These results show that a fraction of missense and synonymous SNPs within EEs significantly disrupt TF binding, with both loss and gain effects, highlighting the potential regulatory consequences of genetic variation within exonic enhancers.
